# Spatiotemporal expression of the zebrafish *pax9* gene that is essential for median fin patterning

**DOI:** 10.64898/2026.09.02.748930

**Authors:** Ziyu Dong, GuangJun Zhang

## Abstract

PAX9 is an evolutionarily conserved paired-box transcription factor that is critical for embryonic development and human diseases. The mouse model has been predominantly used to investigate *Pax9* functions. Zebrafish has emerged as a complementary vertebrate model for various human diseases, including cancers. Until recently, the functions of the zebrafish *pax9* gene in jaw and hematopoiesis have started to be uncovered. However, detailed *pax9* spatiotemporal expression, molecular mechanisms, and potential functions in other zebrafish organs remain largely unknown. With the technical advances in CRISPR-Cas9, non-homologous end joining (NHEJ) has made knockin and knockout a convenient way to examine endogenous gene expression *in vivo* and to generate a loss-of-function allele simultaneously. Here, we first generated *pax9* knockin fish lines by inserting fluorescent proteins at the start of the endogenous *pax9* coding region. Then, we examined *pax9* expression in real time from early embryonic stages through adulthood. Except for previously reported expression domains, we were able to identify *pax9* expression in high resolution in the paired and median fins, where *pax9* marks anterior fin rays. Moreover, our knockin and knockout mutants showed increased fin ray number in median fins, but no evident effect on paired fins in *pax9* null mutants. Thus, PAX9 is critical for median fin patterning in zebrafish.

## INTRODUCTION

Pax genes encode a family of evolutionarily conserved transcription factors defined by the paired DNA-binding domain, a motif shared across metazoans from porifera to mammals ^1^. Generally, there are more *Pax* genes in vertebrates than in invertebrates, mainly due to two consecutive whole-genome duplications (WGDs) ^2,3^. Nine Pax family members (Pax1-Pax9) have been identified and grouped into four subfamilies based on sequence conservation and structural organization in mammals. All of the Pax proteins have a paired domain and a transcription domain (TAD). Group I (*Pax1/9*) is characterized by an octapeptide motif, Group II (*Pax2/5/8*) has a partial homeobox domain in addition to the octapeptide motif, Group III (*Pax3/7*) has a full homeobox domain and an octapeptide motif, and Group IV (*Pax4/6*) has a full homeobox domain but no octapeptide motif ^1,4^. In teleosts, the number of pax genes was further increased by the teleost-specific WGD, which occurred around 240 million years ago ^5–7^. For example, zebrafish have two copies of *pax1* (*pax1a, pax1b*), *pax3* (*pax3a, pax3b*), *pax6* (*pax6a, pax6b*), *pax7* (*pax7a, pax7b*) human orthologues genes according to ZFIN database ^8^.

The *Pax* genes play important roles in vertebrate development and human diseases ^1,4,9^. Vertebrate *Pax1* and *Pax9* share expression in the pharyngeal endoderm, limbs, and sclerotome, a paraxial mesodermal compartment that gives rise to the axial skeleton ^10–14^. Complete loss of *Pax9* in mice leads to newborn death, likely due to a cleft secondary palate. In addition, the *Pax9* null mutants lack pharyngeal derivatives (thymus, parathyroid gland, and ultimobranchial bodies), have defects of neural crest derivatives (cleft palate, tooth, coronoid process, and other craniofacial skeletons), and deformed mesoderm-origin tissue and organs such as preaxial digit duplications of forelimb and hindlimb, extra anterior metatarsals. *Pax9* mutant mice also exhibit cardiovascular defects, including hypoplastic aortas, bicuspid aortic valves, and aberrant pharyngeal arch arteries ^15^. However, as in *Pax1* mutant mice, *Pax9*-deficient mice do not exhibit obvious vertebral defects, whereas compound mutants lack medial sclerotomal derivatives, including vertebral bodies and intervertebral discs, thereby demonstrating functional redundancy and establishing *Pax1/9* as central regulators of vertebral chondrogenesis ^16^. At the molecular level, PAX1 and PAX9 directly activate BAPX1 (NKX3.2) to initiate chondrogenic differentiation of sclerotome-derived progenitors ^17^. Also, PAX1 was reported to compete with SOX9 in regulating aggrecan in murine intervertebral disc annulus fibrosus cells ^18^. In mammals, *Pax9* is also expressed in the dental mesenchyme and marks sites of future tooth formation by integrating FGF8 and BMP4 signaling ^19^. *Pax9*-null mice exhibit arrested tooth development as well as defects in pharyngeal pouch-derived organs ^20^. In humans, heterozygous *PAX9* mutations cause selective tooth agenesis, highlighting a conserved role in odontogenesis ^21^. Beyond these contexts, *Pax9* is also expressed in limb and fin bud mesenchyme, suggesting a potential role in appendage skeleton development ^14^.

Zebrafish has become a prominent animal model of human diseases due to its similarity to vertebrate biology, external embryonic development, and trackable genetics. In zebrafish, embryonic *pax9* expression has been detected in the sclerotome and pharyngeal arches during early somitogenesis by whole-mount in *situ* hybridization (WISH) ^22^. Interestingly, it was found in the dorsolateral part of the somite mesoderm, which is thought to be a unique, newly identified domain of the sclerotome ^22^. This dorsolateral somite domain has recently been proposed to be responsible for dorsal fin formation ^23^. Zebrafish *pax9* null mutants are viable and fertile. However, the homozygous mutant fish lack upper jaws and barbels but have normal pharyngeal teeth ^24^. Another zebrafish study reported that *pax9* is expressed in the caudal hematopoietic tissues, and paired domain insertion allele (*pax9^+23^*) fish showed reduced neutrophils and emergency granulopoiesis failure upon bacterial infection ^24,25^.

Given the evolutionary conservation and pleiotropic effect of the *pax9* gene, its diverse functions in vertebrates remain largely unexplored, especially in teleosts. Part of the reason is the lack of an animal model that enables accurate assessment of *pax9* gene expression in the context of simultaneous loss of function. Recently, various CRISPR-Cas9-mediated knockin strategies based on homology recombination and non-homologous end joining (NHEJ) have been successfully applied to zebrafish ^26–31^. Among them, NHEJ knockin was reported to be effective for introducing fluorescent reporters to visualize endogenous gene expression in zebrafish and medaka ^29–32^. Here, we generated *pax9* knockin and knockout reporter fish lines, which also serve as loss-of-function alleles. We provide a comprehensive spatiotemporal atlas of *pax9* expression from somitogenesis through adulthood. In addition, we report that expanded *pax9* expression domains in the larval stage are associated with extra median fin rays in adults.

## RESULTS

### Efficient NHEJ-mediated knockin *pax9* reporter fish lines

To visualize real-time *pax9* expression in zebrafish, we chose the CRISPR-Cas9-mediated NHEJ strategy and designed two gRNAs around the start codon of the *pax9* locus (**Fig. 1A**). A commonly used 638 bp *hsp70l* enhancer was used to facilitate knockin and robust gene expression ^29^. To compare and ensure high-fidelity *pax9* expression, we also chose a 520 bp *pax9* upstream coding sequence and no enhancer element in the donor plasmid constructs (**Fig. 1B-E**). The knockin efficiency of the injected F_0_ embryos was approximately 3.4-6.0%, based on the pharyngeal FP expression pattern at 6 dpf (days post fertilization) larvae (**Table S1**). However, the F_1_ germline transmission rate from positive F_0_ adult fish ranged from 1.29% to 37.5% (Table S1), indicating a relatively low knockin rate and the need for a large number of screenings. In contrast, positive F_2_ generation fish were generally close to the Mendelian ratio (**Table S1**). Six founder F_1_ fish lines were screened out, and donor constructs and directions were verified by PCR and nanopore sequencing (**Fig. 1F, Fig. S1A**). Consistent with previous reports, both forward and reverse insertion of the constructs with the *hsp70l* enhancer worked. We did not observe a significant difference in fluorescence intensity between the forward and reverse directions of the *hsp70l* enhancer, though variation was noticeable. The *pax9* knockin fish with *hsp70l* enhancer also yielded very close, if not identical, patterns compared to the knockin fish lines with the 520 bp *pax9* upstream sequence and no enhancer at 7 dpf (**Fig. 1G-L**), indicating that the *hsp70l* enhancer does not interrupt the endogenous expression domain or drive ectopic expression in our experiments. Across all six lines, we observed highly consistent expression patterns.

**Figure 1.**
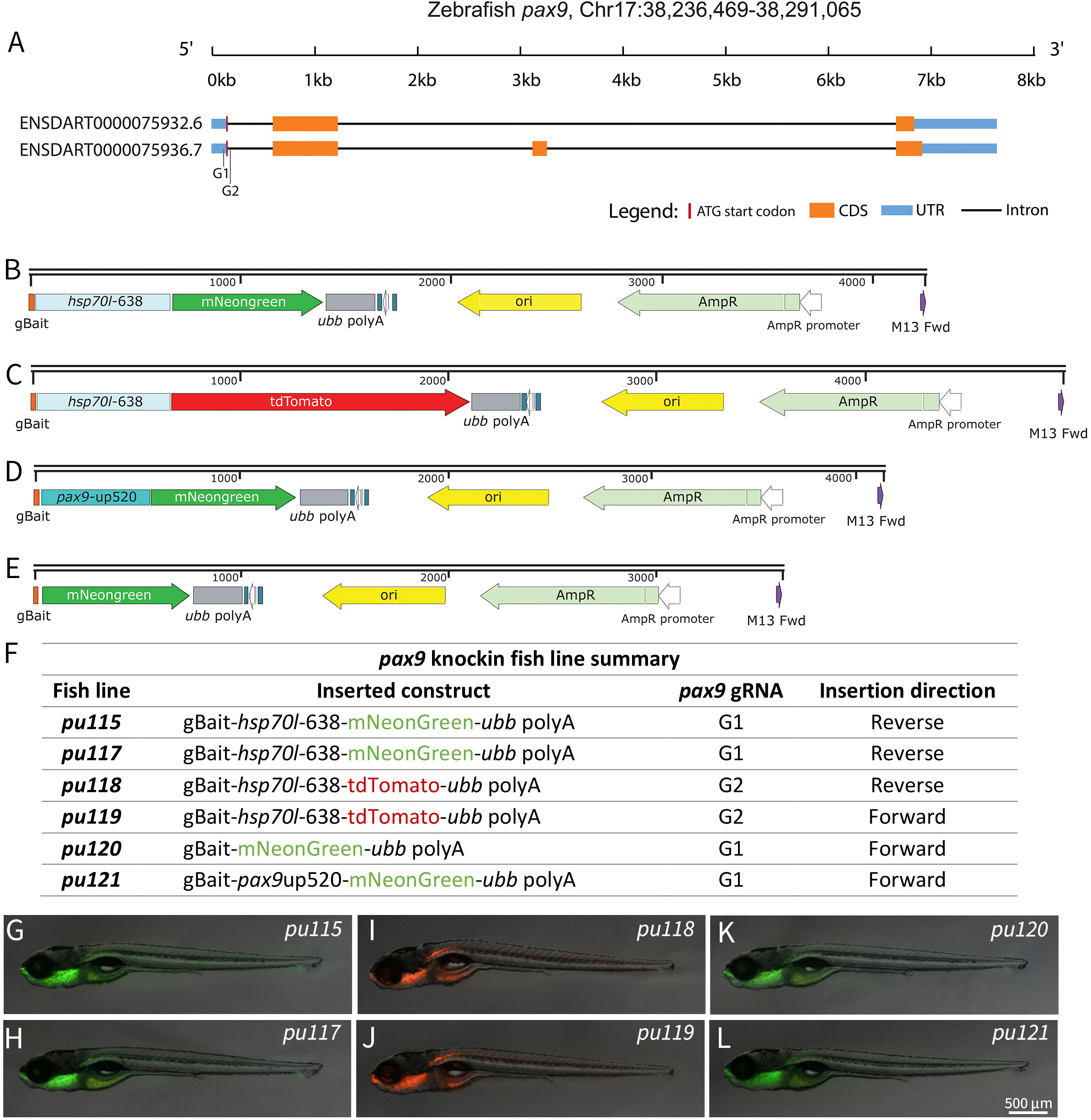
Generation of *pax9* knockin and knockout reporter lines. **A.** Illustration of the zebrafish *pax9* coding region and transcript according to the Ensembl (GRCz11). Two guide RNAs targeting the 5’ region near the start codon and first intron are indicated by black vertical lines. **B-E**. Schematic diagrams of donor plasmids used for *pax9* knockin. **B, C.** Constructs with 638bp *hsp70l* enhancer and mNeonGreen (**B**) and tdTomato (**C**). **D.** Construct with a *pax9* upstream regulatory fragment (*pax9*-up520) driving mNeonGreen. **E.** Construct without an enhancer but with mNeonGreen. All constructs include a gBait sequence, a ubiquitin polyA signal, and a standard plasmid backbone (ori, AmpR). **F**. A summary of knockin fish lines regarding insertional constructs, gRNA, and insertion orientation relative to the *pax9* coding direction. **G-L**. Representative 7 dpf larva whole-mount fluorescence images of the six knockin fish lines.

The 1.5kb *hsp70l* enhancer has been widely used to induce ectopic gene expression in zebrafish ^33–36^. Then we asked whether the 638 bp *hsp70l* enhancer could be adopted for this purpose, in addition to enhancing knockin efficiency. We then performed a brief heat shock (42°C for 1 hour) on 24 hpf (hours post fertilization) *pax9^pu115^* mNeonGreen knockin fish embryos (**Fig. 2A**). Heat-shocked fish embryos showed immediate, universal green fluorescence and remained at least to 48 hpf. In contrast, the untreated fish embryos showed no ectopic fluorescence (**Fig. 2B-J**). Thus, this 638 bp *hsp70l* enhancer not only boosts knockin efficiency, as reported, but can also be activated by heat shock treatment to drive universal gene expression.

**Figure 2.**
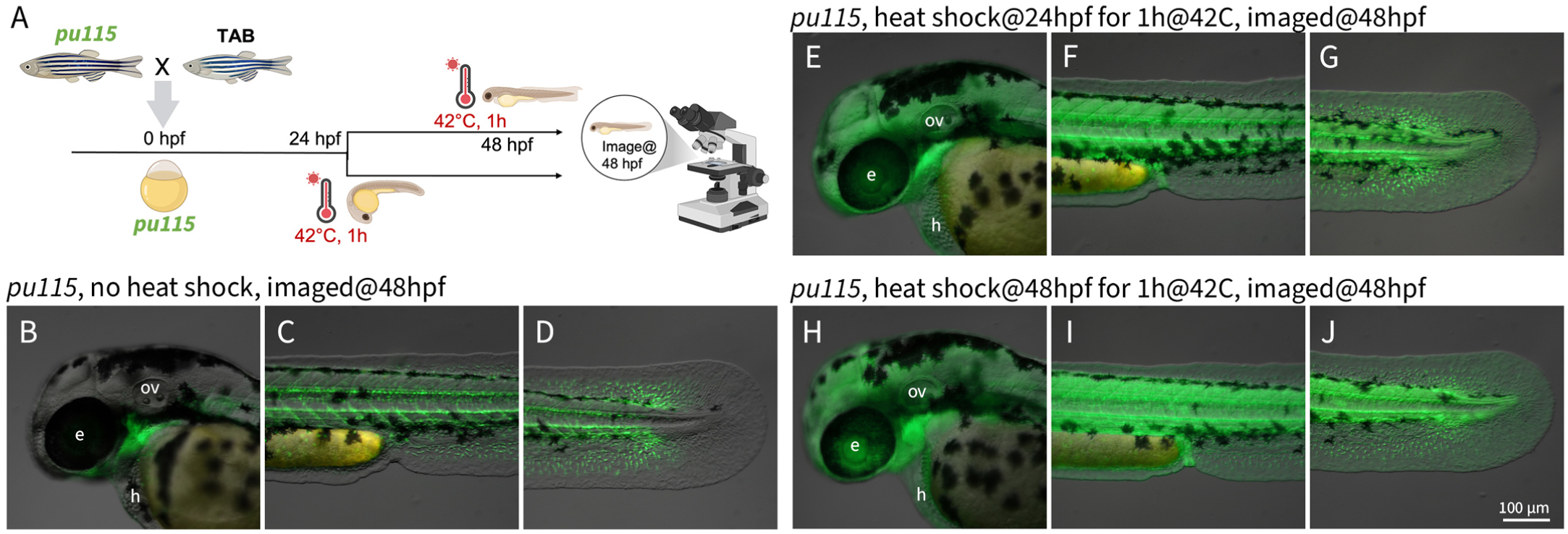
*Pax9* knockin fish fluorescent protein can be activated by a heat shock. **A.** Illustration of heat shock and imaging time frame. Embryos were subjected to heat shock (42°C, 1 hour) at 24 hpf or 48 hpf. All images were taken at 48 hpf. **B-D.** *Pax9^pu115^* embryo mNeonGreen expression in the absence of heat shock. mNeonGreen represents endogenous *pax9* expression. **E-G.** P*ax9^pu115^* embryo heat-shocked at 24 hpf showed increased ubiquitous green fluorescence at 48 hpf. **H-J.** P*ax9^pu115^* embryo heat-shocked at 48 hpf showed immediately widespread green fluorescence. **A**, **E**, **H**, head. **C**, **F**, **I**, trunk. **D**, **G**, **J**. tail. *e*, eye; *h*, heart; *ov*, otic vesicles.

### The *pax9* knockin fish faithfully recapitulate endogenous expression

To evaluate the fidelity of our knockin fish lines, we compared fluorescence to the endogenous *pax9* mRNA distribution using WISH. The *pax9* transcripts were first detectable around the 12-somite stage (12S) and became restricted to the ventromedial somites by WISH (**Fig. 3A-E**). Consistently, the fluorescence of our *pax9^pu115^* fish closely mirrored the mRNA patterns from WISH across all examined stages (**Fig. 3F-J**). All other fish knockin lines showed similar expression patterns at the 22-somite stage, with slightly variable brightness (**Fig. 3K-O**), confirming the high fidelity of these knockin fish for reporting *pax9* endogenous gene expression regardless of insertion orientation. Then, we chose the *pax9^pu115^*(green) and *pax9^pu119^* (red) fish lines for further investigation.

**Figure 3.**
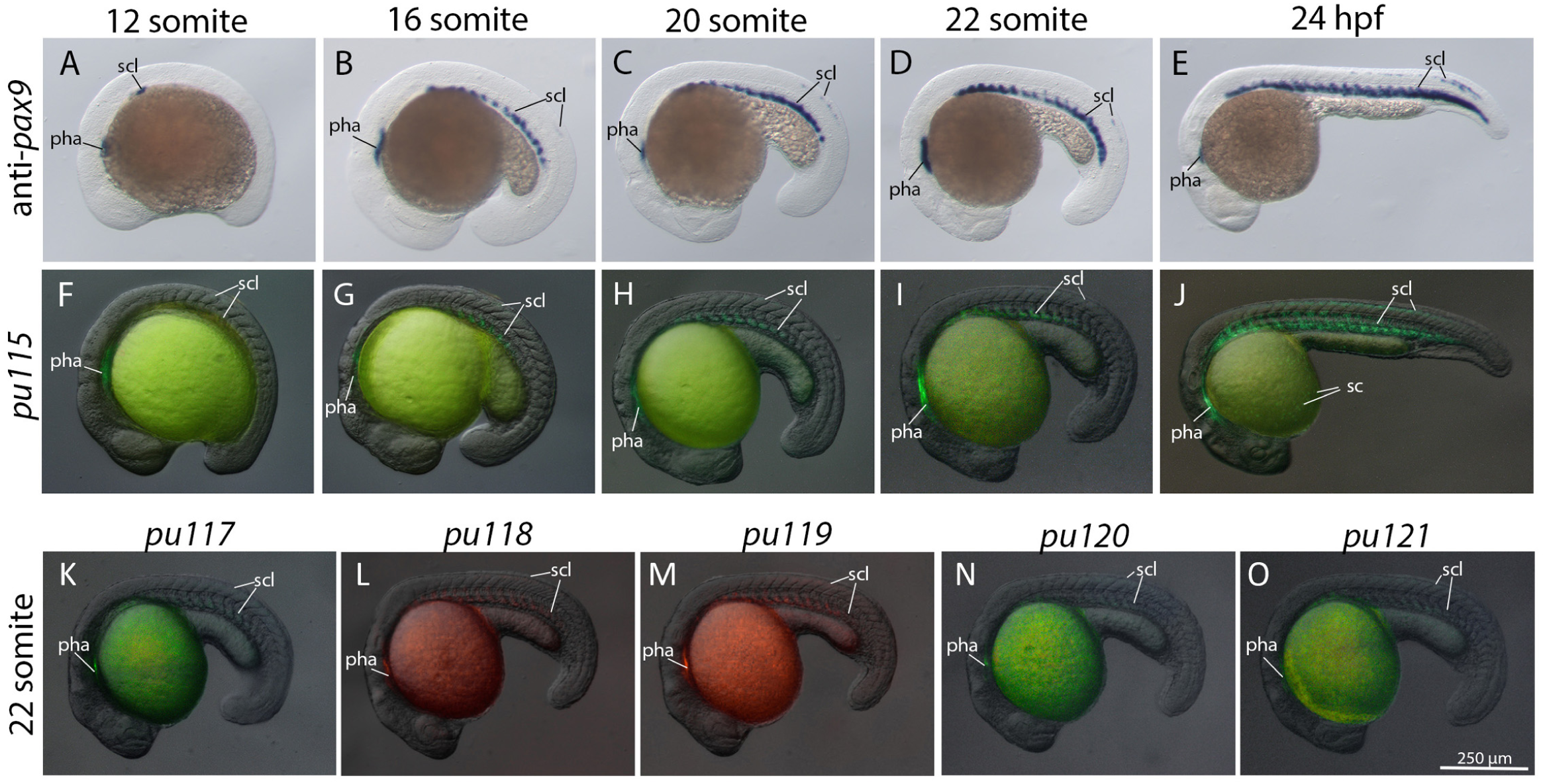
*Pax9* knockin fish mimic endogenous *pax9* gene expression patterns. **A-E.** Endogenous *pax9* mRNA expression by WISH during embryogenesis (12 somite to 24 hpf). Pharyngeal and sclerotome, ventral lateral somites are evident. In addition, *pax9* also labels a zebrafish-specific dorsal lateral domain of mature somites. **F-J.** Corresponding mNeonGreen fluorescence in the *Pax9^pu115^* knockin embryos at the various developmental stages. Some cells on the yolk surface are *pax9*-positive. **K-O.** The 22-somite stage fish embryos across the 6 knockin fish lines showed consistent fluorescence *pax9* expression domains. *pha*, pharyngeal arch; *scl*, sclerotome; *sc*, surface cells.

### The *pax9* gene marks the sclerotome in zebrafish somitogenesis

In *pax9^pu115^* fish embryos, the green fluorescent signal was first detectable at the 12S stage and became restricted to a segmentally repeated ventromedial domain along the anterior-posterior axis during somitogenesis (**Fig. 4A-F**). The anatomical location of this domain corresponds to the sclerotome, the progenitor of the trunk skeleton. Interestingly, *pax9* was limited in the posterior of mature sclerotomes in both WISH and knockin results (**Figs. 3B-O, 4C-F**). To confirm the sclerotome location, we examined the *nkx3.1* gene, an established marker of sclerotome ^22^, and found a well-matched colocalization of *pax9* (GFP) and *nkx3.1* (RFP) expression in the somites at 24 hpf (**Fig. 4G-J**). As in a previous report, we also confirmed the dorsolateral *pax9* expression domain (**Fig. 4C-F**), which is unique to zebrafish ^22^.

**Figure 4.**
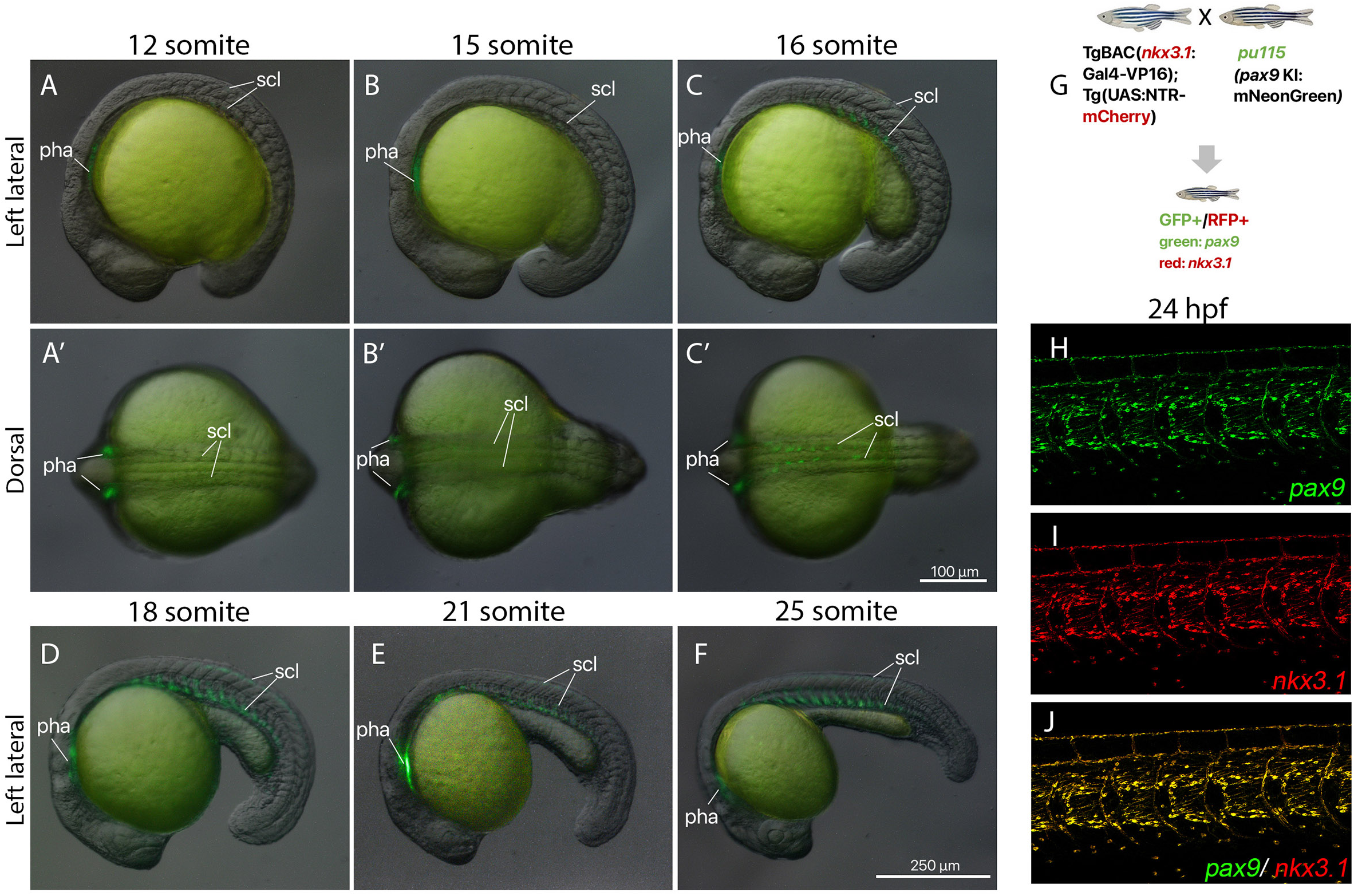
*Pax9* marks the sclerotome during somitogenesis. **A-C.** Lateral views of *pax9^pu115^*knockin embryos at 12, 15, and 16 somite stages. mNeonGreen fluorescence first appears in anterior mature somites and in the pharyngeal arches. **A’-C’**. Dorsal views corresponding to panels **A-C**, highlighting bilateral symmetry of *pax9* expression. **D-F.** Lateral views at the later somitogenesis stages (18, 21, and 25 somites). Sclerotomes are already differentiated in most somites except the newly formed posterior somites close to the tail bud. Zebrafish-specific dorsal lateral sclerotome is evident at these stages. **G.** Schematic of the genetic cross between *pax9^pu115^* and TgBAC(*nkx3.1*:Gal4-VP16);Tg(UAS:NTR-mCherry). **H-J.** A representative sagittal confocal image of embryos showed a well-matched overlap between *pax9* (GFP) and *nkx3.1* (RFP) expression at 24 hpf. *pha*, pharyngeal arch; *scl*, sclerotome.

### *Pax9* expression in paraxial tissue surrounding notochord and developing vertebral columns

At the end of the segmentation stage, *pax9* was expressed in the posterior sclerotome at 24 hpf, and continued in the trunk, around the notochord and myosepta from 2-10 dpf (**Fig. 5A-E**). This axial *pax9* expression domain continued through the larval and juvenile stages, and *pax9* was active in the developing axial skeletons and vertebral columns (**Fig. 5E-H).** In contrast to sclerotomes, *pax9* was highly expressed in the anterior of the vertebral column anlagen. This position exchange is consistent with the previously reported “leaky” resegmentation relationship between somites and vertebral column ^37^. To confirm *pax9* expression in skeletons, we examined the cartilage and bone marker genes *col2a1a* and *sp7/osx*. Indeed, we observed partial overlap between *pax9* and *col2a1a* at 16 dpf and 25 dpf (**Fig. 5I-L**) and between *sp7* at 18 dpf and 25 dpf (**Fig. 5M-P**). This co-expression with either marker only occurs in regions surrounding and interspersed among skeletal elements, suggesting that *pax9-*expressing cells may be closely associated with the differentiation of the two axial skeletal elements.

**Figure 5.**
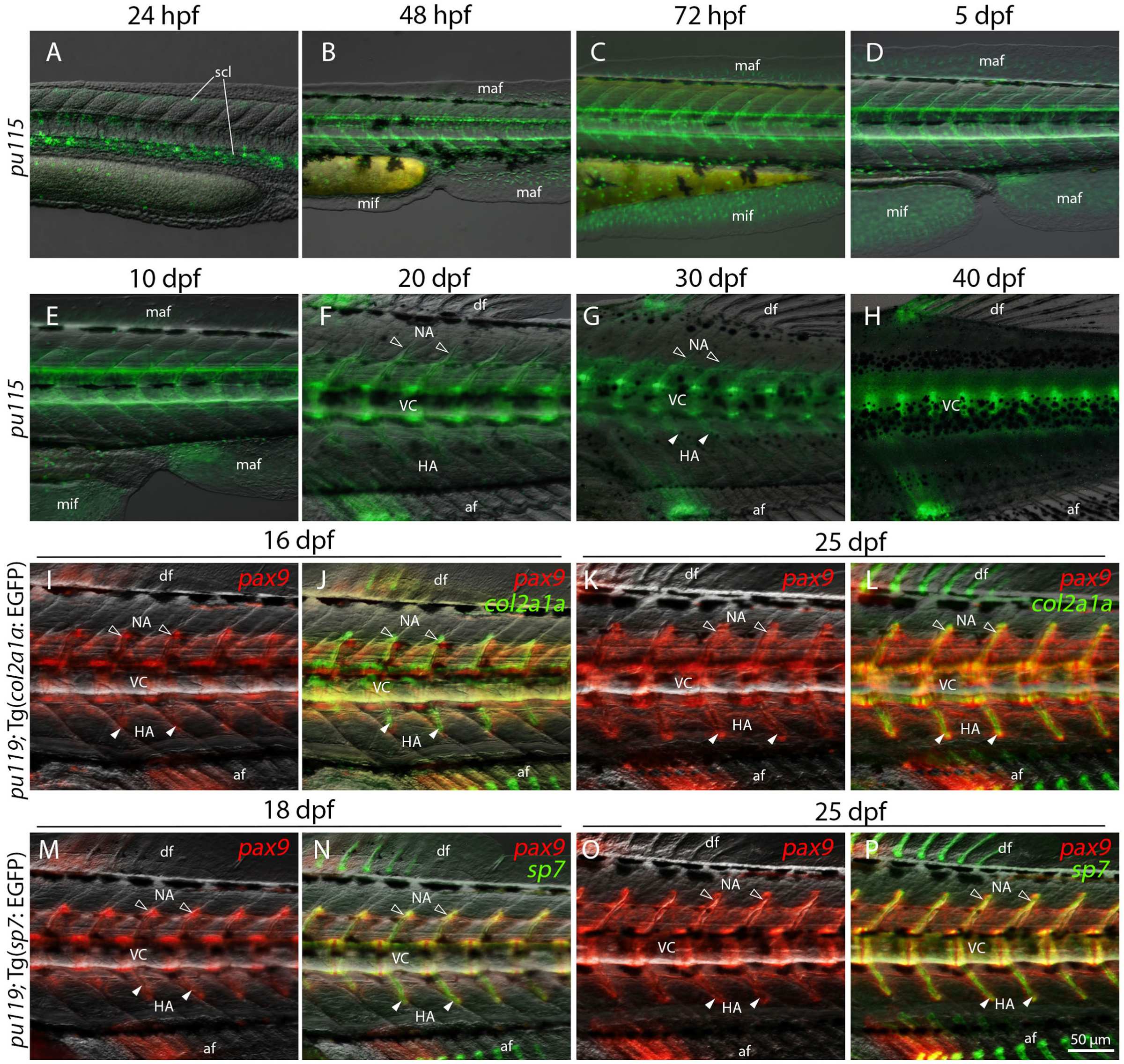
Trunk *Pax9-*expressing cells are associated with axial skeletal structures in zebrafish larvae. **A-H.** Lateral views of trunk regions of *pax9^pu115^* knockin fish from 24 hpf to 40 dpf. The *pax9*-positive cells initially appear in segmentally repeated domains around the notochord during somitogenesis and are maintained through larval and juvenile stages in regions corresponding to the developing vertebral column. **I-L.** Colocalization of *pax9^pu119^* and chondrogenic marker gene, *col2a1a* (Tg(*col2a1a*:EGFP)) at 16 dpf and 25 dpf. **M-P.** Colocalization of *pax9^pu119^* and the osteogenic marker gene, *sp7* (Tg(*sp7*:EGFP)) at 18 dpf and 25 dpf. All images are lateral views of the trunk. *af*, anal fin; *df*, dorsal fin; *HA*, hemal arch (filled white arrowheads); *maf*, major median fin fold; *mif*, minor median fin fold; *NA*, neural arch (open white arrowheads); *scl*, sclerotome; *VC*, vertebral column.

### *Pax9* expression during median fin development

The *pax9* fluorescent signal was detected in the major median fin fold from 2 dpf, but excluded from a distinct posterior domain in the caudal region (**Fig. 6A, B**). This *pax9*-free region was consistently observed and defined a sharp spatial boundary within the caudal fin fold (**Fig. 6A-D**). The *pax9* expression in the minor median fin fold starts at 2-3 dpf (**Fig. 5B, C**). These patterns remain during early larval stages (2-7 dpf). Zebrafish *nkx3.1*-positive sclerotome cells were reported to migrate into the fin folds and become fibroblasts ^38^. Then, we examined the *pax9* and *nkx3.1* expression in the median fin folds and found a closely matched colocalization, indicating that at least some *pax9*-positive sclerotomal cells also migrated and became fin fold fibroblasts.

**Figure 6.**
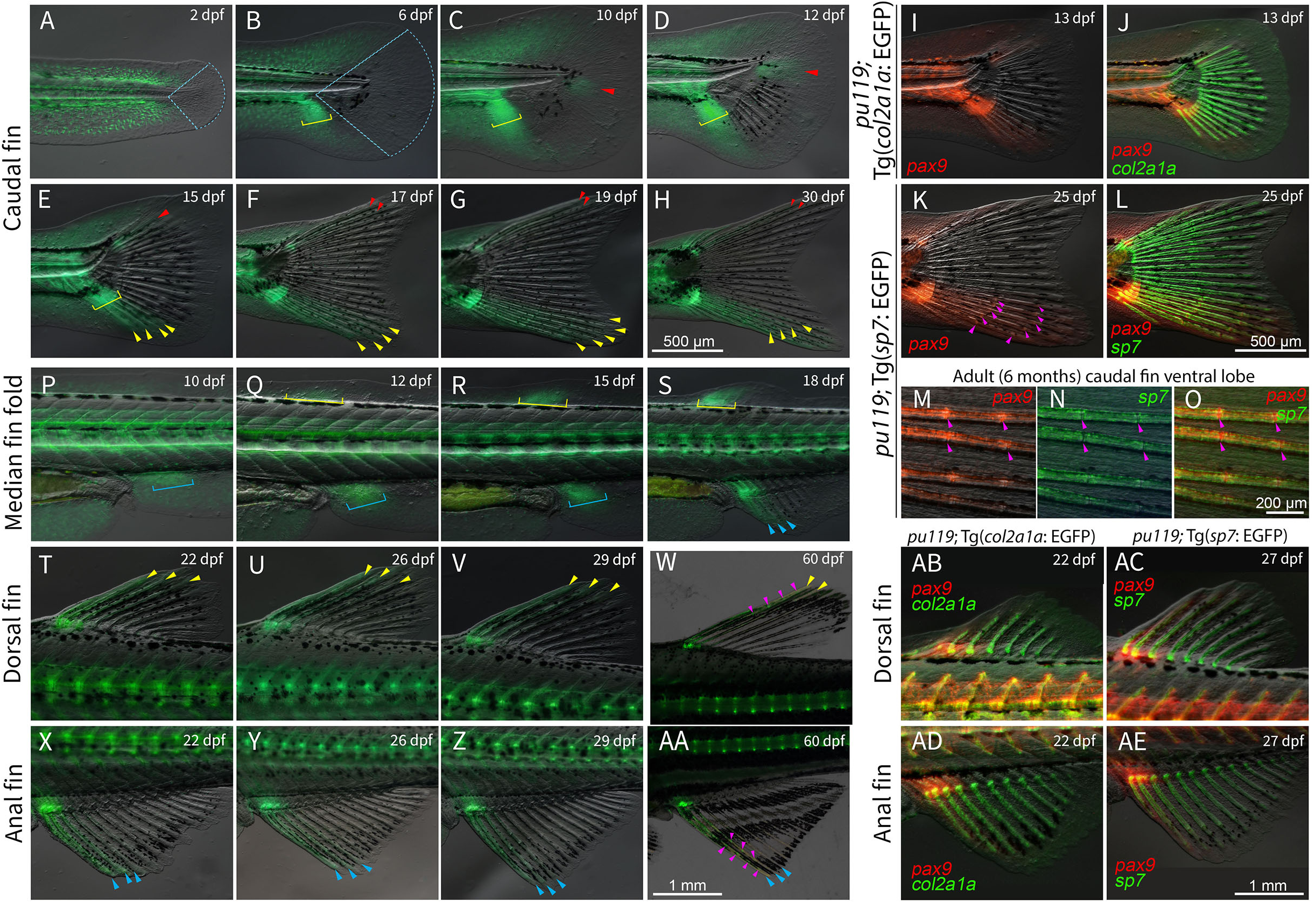
*Pax9* expression delineates mesenchymal condensation and anterior fin skeletals during median fin development. **A-H.** Lateral views of caudal fin from larval to juvenile stages (2-30 dpf) in *pax9^pu115^* knockin fish. *Pax9* is expressed in the major fin fold but forms a *pax9*-free region (blue dotted outline) at 2-6 dpf. *Pax9*-positive mesenchymal condensations (yellow square bracket) are detected in the ventral fin fold and give rise to fin rays (yellow arrowheads). During the notochord flexion period (∼10-12 dpf), a secondary *pax9*-positive domain emerges within the previously pax9-free region (red arrowheads). This secondary *pax9*-positive domain later develops into the two dorsal peripheral principal rays (**D-H**). **I-L.** Colocalization of *pax9* (red) with the cartilage marker, Tg (*col2a1a*: EGFP, green), and with the osteoblast marker, Tg(*sp7*:EGFP, green), in the caudal fin. The *pax9*-positive domains are adjacent to and partially overlap with cartilage and osteoblast populations in the most anterior portion during fin ray development. Magenta arrowheads indicate pax9 expression in fin ray joints. **M-O.** High-resolution views of the adult (6 months) caudal fin ventral lobe with *pax9* (red) and the osteoblast marker, Tg(*sp7*:EGFP) (green). *Pax9* overlaps with *sp7*-positive osteoblasts in peripheral fin ray segments, but not intersegmental joints (yellow arrowheads) that lack *sp7* expression. Magenta arrowheads indicate pax9 expression in fin ray joints. **P-S.** *Pax9* is expressed in the median fin primordia prior to fin ray formation. Yellow square brackets indicate the dorsal fin bud; blue square brackets indicate anal fin primordia. **T-AA.** Dorsal and anal fin development from larval to juvenile stages (22-60 dpf). *pax9* becomes restricted to anterior-most radials and fin rays (yellow and blue arrowheads), while posterior ones remain negative. Magenta arrowheads indicate pax9 expression in fin ray joints. **AB-AE**. Colocalization of *pax9* with the cartilage marker, Tg (*col2a1a*: EGFP, green), and with the osteoblast marker, Tg(*sp7*:EGFP, green), at the most anterior dorsal and anal fins at 22 dpf and 27 dpf, respectively.

Among the zebrafish median fins, the caudal fin develops first. The *pax9* expression increased, and a mesenchymal condensation formed in the ventral caudal fin fold around 4 dpf, and became evident 6-12 dpf (**Fig. 6B-D**). As development proceeded, these condensations gave rise to ventral caudal radials and fin rays, in which *pax9* expression is retained (yellow arrowheads), suggesting a possible role for *pax9* in fin anterior-posterior patterning. When the notochord flexed (10-12 dpf), a secondary *pax9*-positive domain emerged within the previously *pax9*-free region (**Fig. 6C-F**, red arrowheads). This domain was spatially restricted to the dorsal caudal fin region and subsequently contributed to the formation of dorsal peripheral principal rays, reinforcing the role of *pax9* in caudal fin patterning. By juvenile stages (14-30 dpf), *pax9* expression became restricted to the most anterior/ventral caudal fin rays, and its expression in the fin fold recessed (**Fig. 6E-H**). *Pax9* signal was retained in the dorsal-most and ventral-most caudal fin rays at 30 dpf (**Fig. 6H**). There were more *pax9*-labeled rays in the ventral lobe compared to the dorsal lobe, while intermediate rays lacked a detectable mNeonGreen signal. To assess the relationship between *pax9*-expressing cells and skeletal cells during fin development, we examined chondrogenic Tg(*col2a1a:EGFP*) and osteogenic Tg(*sp7:EGFP*) domains. In the caudal fin, *pax9*-positive domains were positioned adjacent to and partially overlapping with *col2a1a*-positive cartilage condensations at early stages (13 dpf) of fin ray formation (**Fig. 6I-J**). At 25 dpf, *pax9* expression showed a close spatial association with *sp7*-positive osteoblasts along developing fin rays (**Fig. 6K-L**). In both cases, *pax9* expression was not uniformly co-localized with either marker, but instead restricted to the regions surrounding or interspersed with skeletal elements. In adult zebrafish, high-resolution images of the ventral lobe showed that *pax9*-positive cells were closely associated with *sp7*-positive osteoblast populations along fin rays (**Fig. 6M-O**). Substantial overlap between *pax9* and *sp7* signals was observed within mineralized ray segments. However, the intersegmental joints expressed *pax9* but not *sp7*. This spatial distinction suggests that *pax9*-expressing populations contribute to multiple fin ray structures, including both mineralized and unmineralized joint tissues.

The anal and dorsal fins start to develop from 10 dpf and 12 dpf, respectively. Before fin ray formation, *pax9* was expressed in anterior mesenchymal condensations within both fin primordia, corresponding to early sites of radials and fin ray formation (**Fig. 6P-S**). As fin rays emerged and elongated, *pax9* expression became restricted to the anterior-most fin rays (**Fig. 6T-AA**). Posterior fin rays did not exhibit detectable reporter expression, indicating a spatial bias along the anterior-posterior axis. This pattern was maintained through later developmental stages, with *pax9* expression confined to a subset of anterior fin rays in both fins (**Fig. 6AB-AE**). Similar to the caudal fin, *pax9* expression was also found at the fin ray joints of the dorsal and anal fins (**Fig. 6W, AA**). Consistent with our observations in the caudal fin, *pax9*-positive domains in dorsal and anal fins were positioned adjacent to and partially overlapping with *col2a1a*-positive and *sp7*-positive skeletal cell populations.

### *Pax9* expression during paired fin development

Zebrafish pectoral fin buds initially formed around 40 hpf, and there was no detectable *pax9* signal within the bud (**Fig. 7A**). As development proceeded (70-85 hpf), *pax9* expression began to emerge within the fin bud around 70 hpf and became clear at 85 hpf (**Fig. 7B, C**). By early larval stages (5 dpf), *pax9* expression was found near the fin base and proximal fin mesenchyme (**Fig. 7D**). As the pectoral fins continued to develop (38-60 dpf), *pax9* became restricted to some anterior fin rays, while the rest of the fin rays remained negative (**Fig. 7E, F**). Zebrafish pelvic fin developed at early juvenile stages, much later than the pectoral fin, and the *pax9* signal was first observed in the anterior base of the fin primordium at 22 dpf (**Fig. 7G**). As pelvic fin rays continued to develop and elongate, *pax9* became confined to the most anterior 2-3 fin rays (**Fig. 7H, I**). In contrast, the remaining rays lacked a detectable signal, similar to pectoral fins. Within these *pax9*-positive anterior fin rays, *pax9* was also expressed highly in the fin ray joints, as in the median fins.

**Figure 7.**
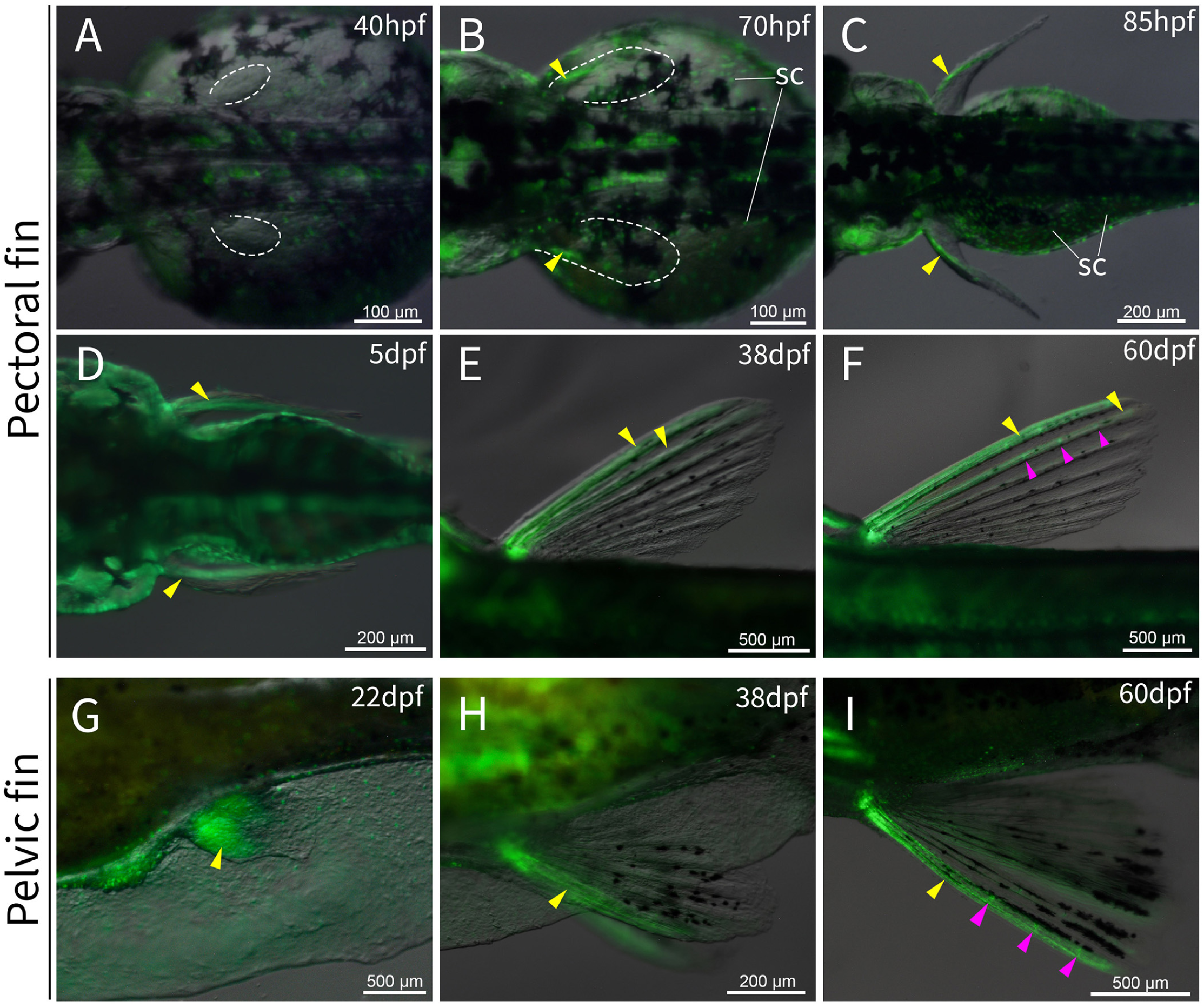
*Pax9* marks anterior paired fins. **A-C.** Dorsal view of early pectoral fin buds (40-85 hpf) of *pax9^pu115^* embryos and larvae. **A**. No *pax9* is detected within the fin bud around 40 hpf. **B**-**C.** *pax9* emerges at the anterior part of the pectoral bud at 70 and 85 hpf. White dashed outlines indicate the pectoral fin buds, and yellow arrowheads mark the onset of *pax9* expression. **D-F.** Pectoral fins of larval to adult fish (5-60 dpf). *pax9* expression is found near the fin base at 5 dpf and becomes restricted to most anterior fin rays and radials between 38 and 60 dpf. **G-I**. Left lateral view of the pelvic fin (22-60 dpf). *pax9* is first observed in the pelvic fin primordia at 22 dpf and later becomes confined to the most anterior fin rays and radials from 38 to 60 dpf. Magenta arrowheads indicate *pax9* expression in fin ray joints. *sc*, surface cells.

### *Pax9* craniofacial expression

*Pax9* was first detected in the pharyngeal region as early as the 12S stage (**Fig. 4A**). At 3 dpf, *pax9* was prominently detected in the pharyngeal arches in the head (**Fig. 8A, D**). By 6 dpf, the *pax9* pharyngeal signal became clear in the pharyngeal arches (**Fig. 8B, E**). Higher-resolution optical sections from confocal z-stacks confirmed that the fluorescence signal is indeed enriched in an outer layer surrounding the pharyngeal arch branches (**Fig. 8B’, E’**). From 3-9 dpf, *pax9* was also expressed in the inner layer of the upper and lower jaw (**Fig. 8A-F**). This jaw expression (inner of maxillary and mandibular) persisted through larva to 20 dpf and adulthood (**Fig. 8G-I**). In addition to craniofacial skeletal regions, *pax9* was found in the nasal region (**Fig. 8G, H**) and maxillary barbels (**Fig. 8I**). The *pax9*-positive cells were located within the central rod of the barbel rather than the surface epithelium (**Fig. 8J**). Interestingly, the barbel inner pole was found to be derived from *sox10*-labeled neural crest ^24^, indicating *pax9*’s pleiotropic function across multiple tissues and organs. On the head surface, we also observed some *pax9*-positive surface cells (**Fig. 8A, D, C, F**), which may be the same cell type found on the yolk surface at 24 hpf (**Fig. 3J**). The identity and developmental origin of these superficial *pax9*-positive cells remain to be determined.

**Figure 8.**
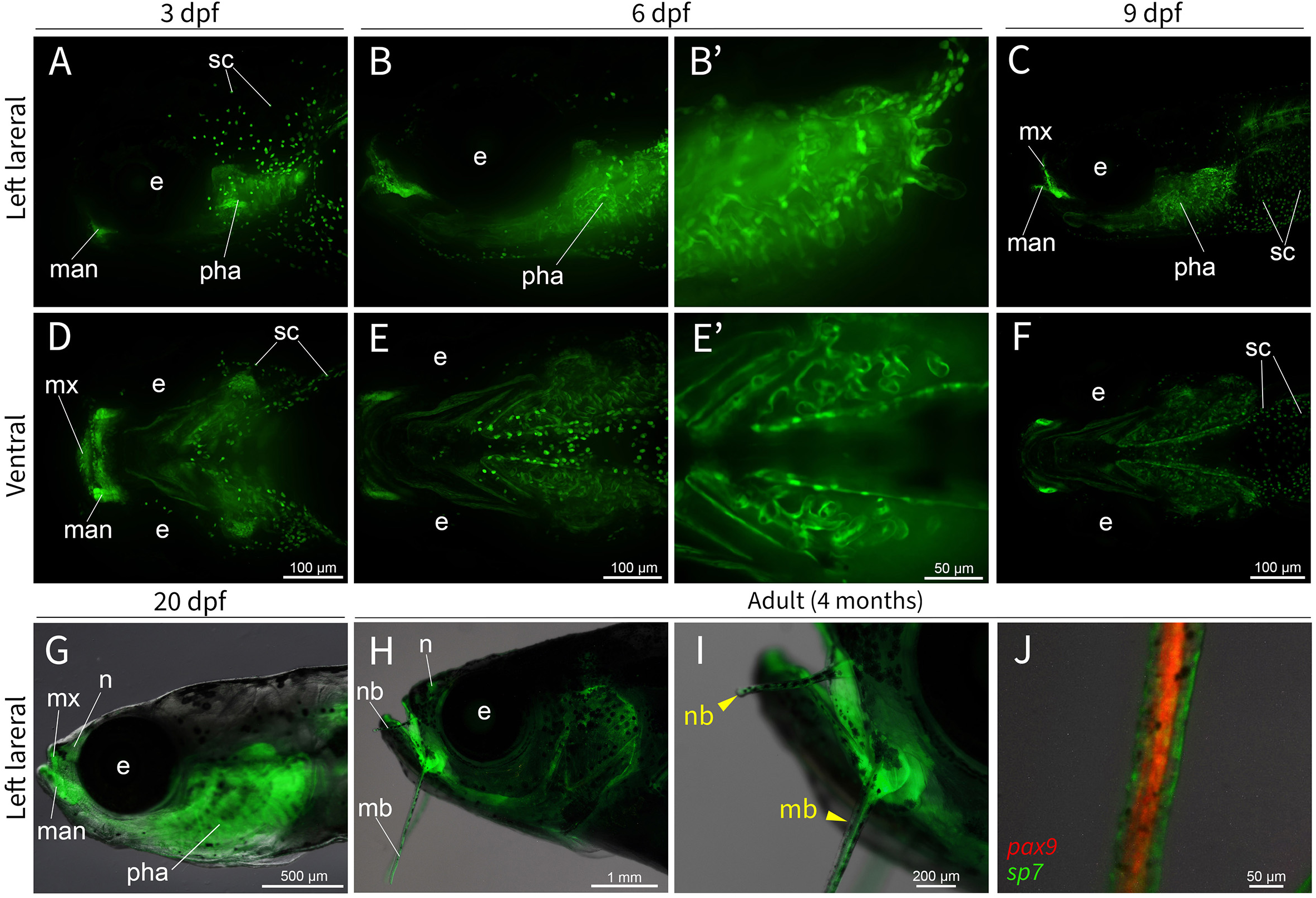
Zebrafish *pax9* expression during craniofacial development. **A-F**. Spinning disk confocal microscopy images. **A**-**C.** Lateral views of *pax9* expression in *pax9^pu115^* larval fish at 3, 6, and 9 dpf. Certain superficial cells are *pax9*-positive (**A**, **C, D, F**). Pharyngeal arches and the inner layer of the jaw are also labeled by *pax9* (**B**, **C**). **D-F.** Ventral views at corresponding stages (3-9 dpf), showing *pax9*’s bilateral expression domains. **B’**, **E’.** Higher-magnification views show the cellular distribution of pax9-positive endodermal cells along the periphery of pharyngeal arch branches. **G.** Lateral view of a 20 dpf *pax9^pu115^* fish larva showing persistent expression in craniofacial structures. Barbels have not developed at this stage. **H-J**. Craniofacial *pax9* expression in adults (4 months). The inner layers of both the upper and lower jaws continue to express *pax9*. **I.** An enlarged view of the mouth region shows *that pax9 is expressed in the* nasal and maxillary barbels (yellow arrowheads). **J**. High-magnification view of the maxillary barbel showing *pax9*-positive cells localized within internal barbel structures, while the outer epithelial layer is largely unlabeled. *e*, eye; *man*, mandible; *mb*, maxillary barbel; *mx,* maxilla; *n*, nasal; *nb*, nasal barbel; *pha*, pharyngeal arch; *sc*, surface cells.

### *Pax9* loss-of-function increases median fin ray number and expands *pax9*-positive ray domains

Because our *pax9* KI fish contains a >4kb plasmid insertion around the *pax9* start codon (**Fig.1A-E**), these fish lines are also *pax9* loss-of-function mutants. Indeed, we observed the typical underdeveloped upper jaw and lack of barbels in our green and red double knockin fish, *pax9^pu115/pu119^* (**Fig. 9B, D**). In addition, double knockin fish showed evident median fin changes, with overlapping GFP and RFP labeling in the *pax9*-positive domains described above for *pax9^pu115^* fish (**Fig. 6**, **Fig. 9B-E**). There was a significant increase in total fin ray number in double knockin fish across all median fins, including caudal, anal, and dorsal fins (**Fig. 9F, H, I**). In contrast, single knockin fish (*pax9^pu115^* or *pax9^pu119^*) were comparable to non-carrier siblings, indicating that a single functional allele is sufficient to maintain normal fin ray number. The double knockin fish caudal fin was asymmetric due to a pronounced expansion in the ventral lobe (**Fig. 9G**). <u>D</u>ouble knockin fish median fins had more FP+ (fluorescent protein-positive) fin rays and an increased ratio of FP+ fin rays to total rays across all median fins, compared to single knockin fish (**Fig. 9J-N**, **Fig. S2A-E**). Thus, our results indicate that a preferential increase in *pax9*-positive rays accompanies the expansion of fin ray number. It is worth noting that the FP+ fin rays are primarily localized to the anterior anal fin. However, additional FP+ rays were observed in the most posterior portion of the anal fin in a subset of double-positive fish (5 out of 18), but not observed in control groups (**Fig. S2F-I**). This phenotypic variation could be due to *pax9* genetic expressivity, as noticed in human tooth agenesis ^39^.

**Figure 9.**
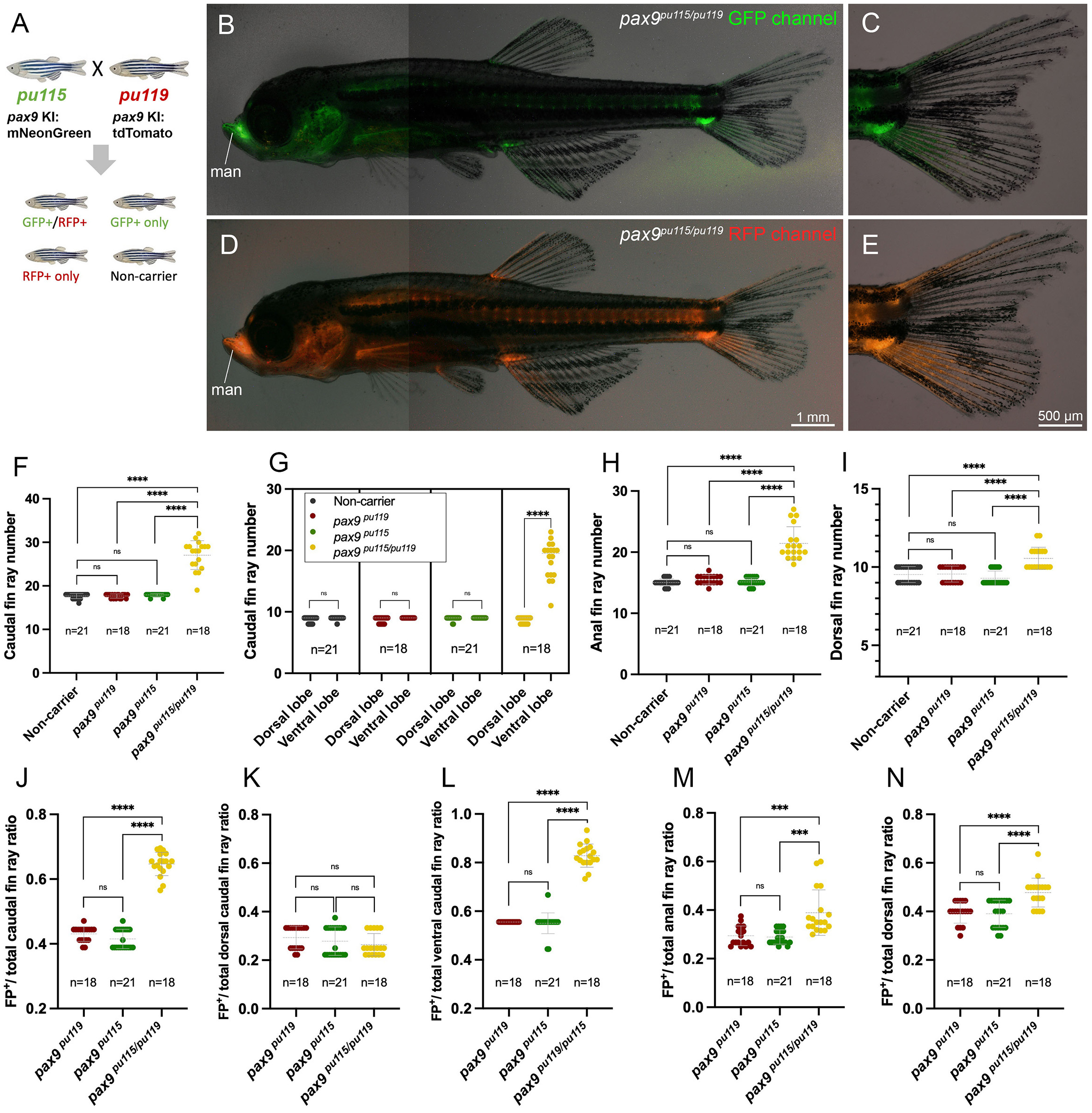
*Pax9* double knockin fish median fins have extra fin rays. **A**. Illustration of generating loss-of-function double knockin (GFP+ and RFP+) fish by crossing *pax9^pu115^*(green) with *pax9^pu119^* (red). **B-E.** Representative double-positive adult fish (30 dpf), which express both green **(B, C)** and red fluorescent **(D, E)** proteins at the same locations. **F, H, I.** Quantification of total fin ray number in caudal, anal, and dorsal fins across all genotypes (non-carrier, *pax9^pu119^, pax9^pu115^, and pax9^pu115/pu119^*). **G.** Quantification of dorsal and ventral caudal fin ray numbers. Fin ray number increases in the ventral caudal lobe, but not the dorsal lobe. **J-N.** Quantification of fluorescent protein-positive (FP+) fin rays over the total fin ray ratio in caudal, anal, and dorsal fins. *ns*, not significant. *** indicated significance with p< 0.001. **** indicated significance with p< 0.0001. *man*, mandible.

### Trans-heterozygous and homozygous mutants recapitulate median fin phenotypes

To validate our findings in the double knockin fish, we then generated *pax9* loss-of-function knockout mutant fish lines, *pu116* and *pu122*, using a gRNA targeting the 2^nd^ and 4^th^ exons (**Fig. S1B**). Both mutants are predicted to produce truncated proteins due to their indel mutations. Then we examined trans-heterozygous mutants by crossing *pax9^pu115^* with *pax9^pu116^*. Trans-heterozygous mutants (*pax9^pu115/pu116^*) recapitulated the phenotypes observed in *pax9^pu115/pu119^* double knockin fish, including increased fin-ray number in the caudal, dorsal, and anal fins (**Fig. 10A-D**). This confirms that the observed median phenotypes are attributable to loss of *pax9* function. Interestingly, the enlarged fin partially resembles the mutants of combinational *hox* genes (*hoxc12a;hoxc13a;hoxc12b;hoxc13b*) ^40^ and *hhip* ^41^, the antagonist of Hedgehog signaling, suggesting PAX9 might interact with the posterior HOX proteins and members of the Hedgehog signaling pathway.

**Figure 10.**
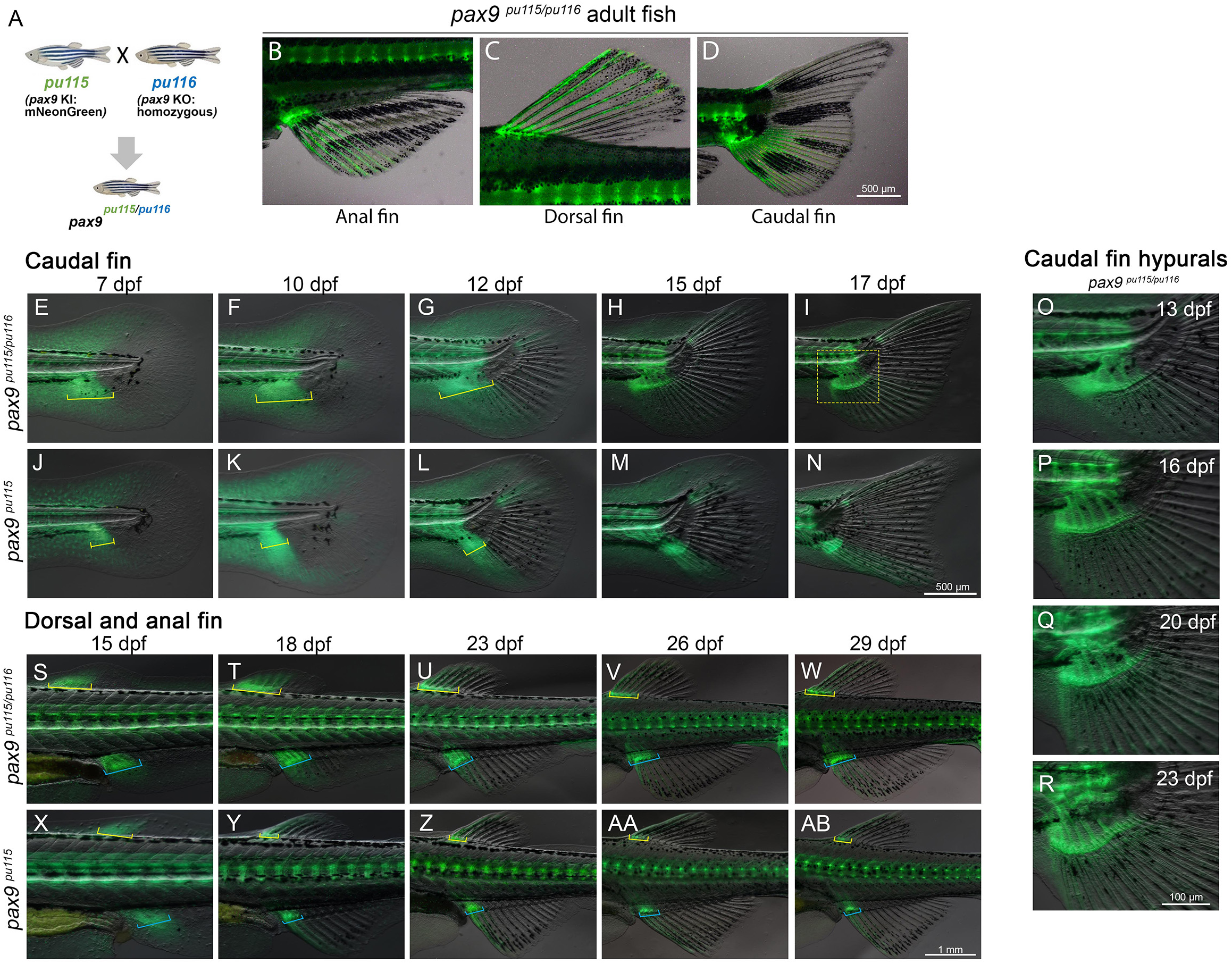
Trans-heterozygous *pax9* mutants develop an enlarged mesenchymal condensation during median fin development. **A.** Illustration of the trans-heterozygous strategy by crossing *pax9^pu115^* knockin fish with the *pax9^pu116^*knockout fish. **B-D.** Representative adult fins (anal, dorsal, caudal) in trans-heterozygotes, *pax9^pu115/pu116^* fish, with extra fin rays. **E-N.** Caudal fin developmental process (7-17 dpf) in trans-heterozygotes (**E**-**I**) and *pax9^pu116^* control fish (**J**-**N)**. Yellow square brackets indicate the mesenchymal condensations. **O-R**. Higher-magnification views of the caudal fin hypural region. The number of hypural elements increases in trans-heterozygous fish. **S-AB.** Developmental process of dorsal and anal fins (15-29 dpf). Fin ray number increases in trans-heterozygous fish *pax9^pu115/pu116^* (**S**-**W**), compared to control fish *pax9^pu115^* (**X**-**AB**). Yellow and blue square brackets indicate the mesenchymal condensations of dorsal and anal fins.

To understand the developmental process of these extra median rays, we tracked the *pax9* signal in median fins across developmental stages in trans-heterozygous and control fish. We found that *pax9-expressing* mesenchymal condensations expanded in mutants compared to controls (**Fig. 10E-N**). This expansion is particularly evident in the hypural region, where increased numbers of hypural elements were observed between 13-23 dpf (**Fig. 10O-R**). As fish developed, the *pax9*+ expanded condensations gave rise to additional internal skeletons, each associated with more fin rays. Similar trends were observed in dorsal and anal fins, where expanded *pax9*-positive mesenchymal domains preceded extra radials and fin rays (**Fig. 10S-AB**). Together, these results support the idea that loss of *pax9* function leads to expansion of *pax9*-positive mesenchymal domains in median fin primordia, which differentiate into more radials. The increased fin ray numbers seem secondary to the extra endoskeletons, hypurals, and radials.

Next, we further validated the median fin phenotypes by examining homozygous *pax9* knockout mutant fish, *pax9^pu116/pu116^,* using alcian blue and alizarin red staining. We confirmed that enlarged mesenchymal condensation adopted a chondrogenic fate in the caudal fin primordia, and more hypural elements formed in mutants (**Fig. 11A-D**). Consistent with our transheterozygous mutants, more ventral caudal fin rays are associated with the extra hypural elements (**Fig.11E-F**). We also observed the lack of the upper jaw and maxillary barbels (**Fig. 11G-J**), as previously reported ^24^. In addition, the ventral caudal fin ray number in *pax9^pu116/pu116^* and *pax9^pu122/pu122^* fish increased similarly to those observed in double knockin fish (*pax9^pu115/pu119^*) and trans-heterozygous fish (*pax9^pu115/pu116^*) (**Fig. 11K-U**), in contrast to the unchanged pectoral and pelvic fins (**Fig. S2J-K**). Thus, our results confirm that increased fin rays are robust in these different *pax9* loss-of-function mutants.

**Figure 11.**
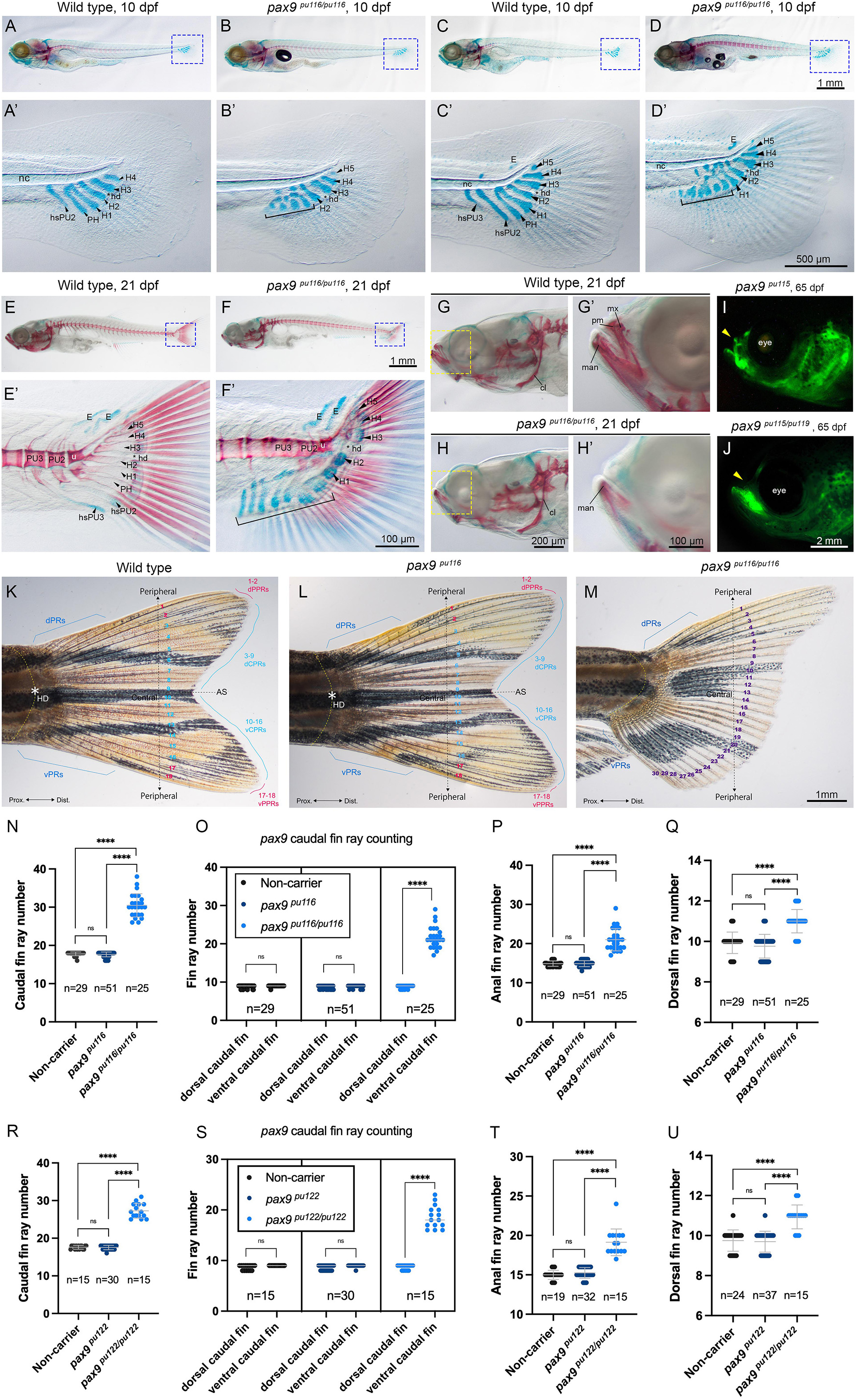
*Pax9* null mutants recapitulate the median fin phenotypes observed in trans-heterozygotes. **A-H.** Left lateral view of wildtype (**A**, **C**, **E**, **G**) and *pax9* knockout *pax9^pu116/pu116^*(**B**, **D**, **F**, **H**) larval and juvenile zebrafish stained with alcian blue and alizarin red. Cartilages are stained blue, and bones are stained red. **A’-H’**. Higher magnification of the dashed-boxed regions in panels **A**-**H.** Normal hypural elements are numbered, and extra ones are indicated by black square brackets. Bones of the upper jaw are missing in the null mutant compared to the wildtype (**G**, **H**, **G’**, **H’**). **I**-**J**. Fluorescence imaging of the head region shows the lack of barbels (yellow arrowheads) in the null mutant. **K**-**M**. Representative images of fish tails from wildtype (**K**), heterozygote (**L**), and homozygote (**M**). Caudal fin rays are numbered dorsoventrally according to their position relative to the midline landmark, the hypural diastema. **N**-**Q**. Quantification of total fin ray number in *pax9^pu116^* mutant median fins. **R-U**. Quantification of total fin ray number in *pax9^pu122^* mutant median fins. Increased fin rays are located on the ventral side of caudal fins (**O** and **S**). *ns*, not significant. **** indicates significance with p< 0.0001. Asterisk, hypural diastema. *AS*, axis of external dorsoventral symmetry; *cl*, cleithrum; *dCPRs*, dorsal central principal rays; *dPPRs*, dorsal peripheral principal rays; *dPRs*, dorsal procurrent rays; *E*, epural; *eye*, eye; *H*, hypural; *hd*, hypural diastema; *hsPU*, haemal spines of the preural centra; *man*, mandible; *mx*, maxillary; *nc*, notochord; *PH*, parhypural; *pm*, premaxillary; *PU*, preural centra; *u*, urostyle; *vCPRs*, ventral central principal rays; *vPPRs*, ventral peripheral principal rays; *vPRs*, ventral procurrent rays.

## DISCUSSION

The *pax9* gene plays a critical role in vertebrate embryonic development and human diseases. Using the NHEJ-based CRISPR method, we established 6 zebrafish *pax9* knockin-knockout fluorescent fish lines that faithfully recapitulate endogenous *pax9* gene expression. Zebrafish *pax9* gene expression was systematically examined from live early embryos through adulthood. In addition, loss of *pax9* leads to an increase in the number of fin rays of the median fins.

CRISPR knockin fish offer advantages over traditional mRNA detection by WISH, which is static and has limited riboprobe penetration, especially in zebrafish larvae and adults. Similarly, Immunohistochemistry is commonly used to assess protein expression, but it cannot be used for real-time tracking. The employment of the 638 bp *hsp70l* enhancer not only improves KI efficiency due to no preferential insertional orientation but also boosts the fluorescent protein expression. In addition, the knockin fish can be used to temporally overexpress any gene of interest via heat shock when the gene is co-expressed with FP. In our hands, the efficiency of this knockin-knockout approach is acceptable but not high. Moreover, this approach could be easily adopted to Gal4, Cre recombinase, or other proteins of interest for various functional studies.

*Pax9* expression in pharyngeal arches and the sclerotome is conserved across vertebrates, as mouse, zebrafish, skate, and lamprey share these two expression domains ^20,42,43^. This phenomenon indicated that the origin of *pax1/9* after WGDs could be critical for vertebrate-defining characters, such as the jaw and vertebral column ^43,44^. In addition to many shared gene expressions (e.g., *shh*, *fgf8*, etc.), it has long been noticed that the gill arch branchial ray pattern is also similar to that of fin/limb skeletal branches. This is known as the pharyngeal origin hypothesis of the paired fins/limbs ^45–48^. In our *pax9* KI fish, pharyngeal arch expression was detected during somitogenesis and persisted in the arches’ endoderm in larval fish. This is consistent with *Pax9* activity in the pharyngeal pouch-associated mesenchyme in mouse and skate ^20,42^. Expression in the upper jaw and craniofacial mesenchyme of adult zebrafish aligns with craniofacial defects reported in *pax9* mutant fish ^24^. Our *pax9* null mutant results also confirmed these cranial phenotypes. Zebrafish *pax9* sclerotome expression is initiated during somitogenesis and maintained through vertebral column formation. This is similar to the well-established role of *Pax9* in the paraxial mesenchyme in mice ^10,16^. However, the *pax9* null mutant fish do not show evident axial skeletal defects, likely due to the overlapping function of *pax1a* and *pax1b*. The future examination of double mutants of *pax1* and *pax9* may clarify this.

Appendages play important roles in vertebrate adaptation. Median fins appeared in primitive vertebrates before paired fins. The median fins may have evolved from the axial skeleton by co-opting the genetic program of the paraxial mesoderm, specifically the somites ^49^. In contrast, the paired fins may have been co-opted from the pharyngeal arch or median fin. The fin-fold theory proposed that paired fins evolved through the duplication and modification of the median fin fold along the midline. Indeed, more and more genetic and cellular signals and pathways are conserved between paired and median fins ^7,50,51^. It is worth noting that a recent study reported that the unpaired pre-anal fin fold in zebrafish can be derived from the lateral plate mesoderm, based on a few marker genes ^52^. The unpaired pre-anal fin fold may suggest an intermediate structure between median and paired fins ^52^. Although no adult fin skeletons in living vertebrates are related to this transient embryonic structure, this study provides new developmental evidence for the finfold hypothesis. Based on current available experimental data and palaeontological evidence, the origins of paired fins remain debated. Accordingly, we found both similarities and differences in *pax9* expression between paired fins and median fins.

Zebrafish *pax9* is expressed in the anterior mesenchyme of both paired and median fin primordia during the embryonic and larval stages. With the start of mesenchymal condensation, *pax9* is activated in the anterior fin buds, and its expression remains into adulthood. The *pax9* expression of the anterior fin rays is robust, even in fin regeneration, and this pattern is similar to *alx4a* ^53,54^. Since *pax9*’s anterior fin bud expression has also been reported in Xiphophorus and sharks ^48,55^, it may indicate that *pax9*’s role in anterior-posterior fin patterning is evolutionarily conserved. Our collaborative work also supported this patterning function ^56^. Indeed, the shift of this anterior– posterior axis was proposed to be related to the fin-to-limb transition ^48^. One difference is that *pax9* is not found in the paired fin folds but is expressed in the median fin folds before fin primordia emerge. These *pax9*-expressing cells in the median fin fold are likely the cells that migrate from the sclerotome. We have noticed that *pax9* and *nkx3.1* expression overlap well, and *nkx3.1*-expressing fibroblast cells are known to originate from the sclerotome ^38^. This notion is consistent with the somite origin of median fins ^49,57–59^ and is also supported by our collaborative scRNA-seq study on loss-of-function median fin morphology ^56^. Consistently, a recent preliminary lineage-tracing study has shown that dorsal sclerotome cells contribute to the dorsal and anal fins, and that sclerotome-deficient smoothback (*smb*) mutant lacks these structures ^23^. Another major difference between paired and median fins is their response to the *pax9* loss. We do not observe evident morphological changes in the paired fins. In contrast, all median fins show increased fin-ray numbers, though the severity may vary across the genetic backgrounds of different zebrafish strains ^56^. This median-fin-only phenotype could be explained, at least partially, by the paired fins originating from lateral plate mesoderm, while the median fin develops from somites. Different developmental origins may lead to distinct cellular and genetic programs, such as SHH signaling, which is present in the zone of polarizing activity (ZPA) of paired fins, but ZPA is absent in median fins. Our findings on *pax9* expression and function could provide additional insight into the evolutionary origin of fins and fin-ray patterning in teleosts. In addition, due to pleiotropy and genetic expressivity, a slight shift in the *pax9* gene’s expression domain and/or levels across vertebrate species could be critical for adaptation to their environments by fine-tuning jaw, barbel, and median fin shapes.

Loss of *pax9* leads to extra skeletal elements in mouse limbs, including duplicated preaxial digits and additional anterior metatarsals. Zebrafish null mutants exhibit expanded *pax9*-positive mesenchymal condensations at early developmental stages and increased median radials and fin ray numbers in the posterior portion of adult fish. In addition, colocalization with Tg(*col2a1a*: EGFP) and Tg(*sp7*: EGFP) in the axial skeleton (**Fig. 5)** supports its association with skeletogenesis. These mouse and fish data suggest that *Pax9* may negatively regulate skeletal formation or differentiation. When *pax9* is inactivated, more mesenchymal progenitors are committed to skeletal formation. This model is consistent with the established role of PAX1 and PAX9 in activating BAPX1/NKX3.2 to promote chondrogenic differentiation of sclerotome cells in mice ^17^. Furthermore, PAX1/9 has been reported to compete with SOX9, a master chondrogenic regulator, at the aggrecan promoter in cartilage matrix ^18^. Thus, it is also possible that the *pax9* null mutants generated their expanded median fin skeletons simply through an enhanced skeletogenesis at the anterior of the median fins, as illustrated in our model (**Fig. 12**). The presence of ZPA in paired fins, especially the SHH signaling, might bypass the *pax9* inhibitory effect on skeletogenesis through *pax9* downstream genes. So the *pax9* null mutants show phenotypes only in median fins, not in paired fins. This model could be complementary to the sclerotome deployment model for the median fin patterning ^56^. Future RNA-seq on *pax9* mutants could help identify downstream mechanisms of skeletogenesis. Long-term lineage-tracing studies (*e.g.*, CRE-LOXP) are also critical for elucidating *pax9* functions across diverse tissues and organs.

**Figure 12.**
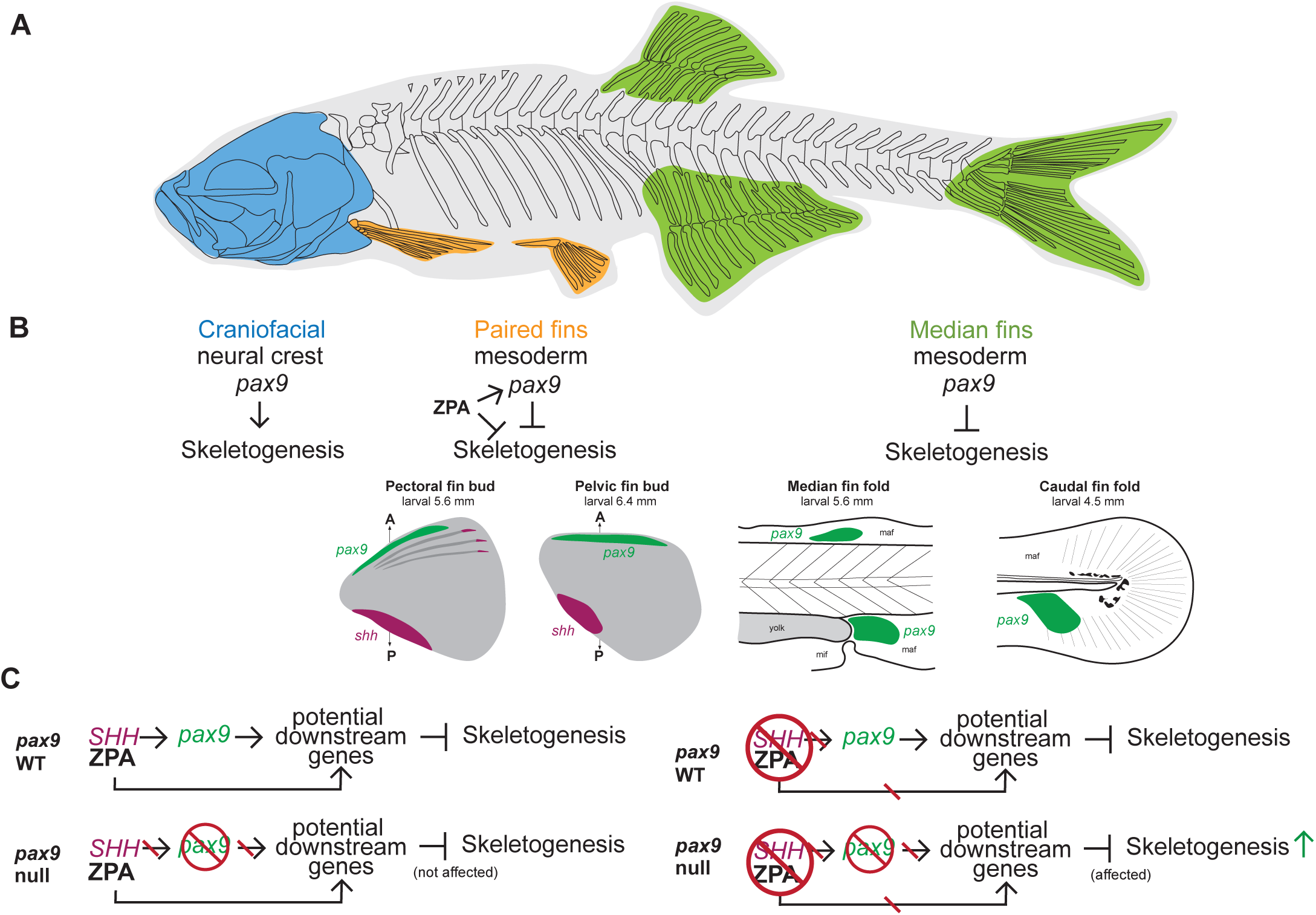
Model of *pax9*’s functions in zebrafish skeletogenesis. **A**. Illustration of adult zebrafish skeletons: craniofacial skeletons derived from neural crest (blue); paired (orange) and median (green) fin skeletons originated from lateral plate and paraxial mesoderm, respectively. **B**. *Pax9* promotes skeletogenesis of the zebrafish maxillary jaw (neural crest origin), while it inhibits skeletogenesis of paired and median fins. **C**. Hypothetical mechanism of *pax9* in paired and median fins. The ZPA of the paired fins, a source of SHH signaling, might bypass the *pax9*’s skeletal inhibitory effects through directly activating potential downstream genes of *pax9*. Red reverse slashes without or within red circles indicate loss or absence. Arrows and ⊣ indicate promoting and inhibiting effects, respectively. *maf*, major median fin fold; *mif*, minor median fin fold.

## MATERIALS AND METHODS

### Zebrafish husbandry and zebrafish fish lines

Zebrafish were raised and maintained in accordance with AAALAC-approved standards at the Purdue animal housing facility, following protocols approved by the Purdue Animal Care and Use Committee (PACUC, #1210000750). All the zebrafish experiments were carried out in a wildtype TAB (AB/Tübingen, RRID: ZIRC_ZL1) background ^60,61^. Zebrafish were maintained according to the Zebrafish Book ^62^, and all zebrafish embryos and larvae were staged according to the Kimmel and Parichy staging guides, respectively ^63,64^. Published transgenic lines used in this study include *Tg(sp7*: *EGFP*)*b1212* ^65^, Tg(UAS: NTR-mCherry)^c264^ ^66^, TgBAC(*nkx3.1*: Gal4)^ca101^ ^22^, Tg(*col2a1*: GFP) ^67^.

### Generation of *pax9* knockin lines

The *pax9* CRISPR-Cas9 knockin fish lines (*pu115, pu117-121*) were generated using an NHEJ-based strategy modified from Kimura et al. ^29^. A guide RNA (gRNA) targeting exon 1 of *pax9* (G1: 5’-GACTCGGAACAGGTCAGAAT-3’) and another gRNA targeting the first intron (G2: 5’-ATGCATGCCCGCTCCCATGT-3’) were used to mediate targeted integration. Donor plasmids were from those of Dr. Mathew Harris Lab and modified by truncating the 1.5kb *hsp70l* enhancer to 638 bp. A donor plasmid containing the gBait sequence (5’-GGCGAGGGCGATGCCACCTA-3’), a 638 bp *hsp70l* enhancer, and the fluorescent reporter genes, mNeonGreen and tdTomato, followed by a polyadenylation signal, was co-injected with Cas9 protein (PNABio, CP01-50), the *pax9*-targeting gRNA, and the gBait gRNA into one-cell stage embryos. For *hsp70l* enhancer control, 520 bp of *pax9* upstream of the start codon ATG, or no enhancer, was added before the fluorescent reporter. Founder fish were identified by fluorescence screening of F_0_ and F_1_ progeny at 6 dpf, and F_2_ stable lines were established by outcrossing F_0_ and F_1_ to TAB fish. Nanopore sequencing was used to confirm targeted integration at the expected *pax9* locus. A detailed step-by-step protocol is available in the supplementary material (Supplementary Protocol).

### Generation of pax9 knockout allele

The *pax9* CRISPR-Cas9 knockout lines (*pu116 and pu122*) were generated using four gRNAs targeting exon 2 and exon 4 using our published CRISPR mutation methods ^68^. CR1: 5’-CCAACTCTACTATCCTGAGCCGG-3’, CR2: 5’-AATCGGCGGCAGTAAACCGAGGG-3’, CR3: 5’-GGGGTACAGCGGCACCACGT-3’, and CR4: 5’-CGAATGGACGGAGTCTCGTG-3’. The resulting mutants were cloned and confirmed by Sanger sequencing. The *pu116* carries a 77 bp deletion (ACGGCTGCGTGAGCAAGATTCTGGCTCGGTACAACGAAACCGGCTCAATACTTCCCGGTGCAATCGGCGGCAGTAAA) and a 43-bp insertion (CCCAAACGTAGTCAAGCACATTAGGACTTACAAGCAACGGGAC) in exon 2, an indel (Deletion of CA and insertion of TATG), and a 10 bp deletion (CCACACGAGA) in exon 4. The *pu122* carries an 18 bp deletion (GCGGCAGTAAACCGAGGG) in exon 2 and a 120 bp deletion (GCACCACGTCGGCCTATGTGACCGGCCCCACATGGCAGCCGCCCAGTGGCAGTGCTCTC TCTCCCCACAGCTGTGACATTAGCAGCCCGCTGGCATTCAAGAGCATGAGTGCCACACGAG) and 1 bp insertion (T). Both are predicted null mutants with truncated proteins due to reading frame shifts (**Fig. S1B**). The *pax9^pu116^* fish were genotyped using PCR with primers specific to the mutant allele within exon 2: 5’-GCATAACTAGAGCCAGCCTTTGG-3’ and 5’-GCGGAGAATCCTACTAATTGAGCTG-3’. Wildtype allele yields a 390-bp band, and the mutant allele produces a 356-bp band. The *pax9^pu122^* fish were genotyped using PCR with primers specific to the mutant allele within exon 4: 5’-CGAATGGATTGCCCACAGTGAAC-3’ and 5’-CATGTAGAACGAGCCAGACTGTTG-3’. Wildtype allele yields a 307-bp band, and the mutant allele produces a 188-bp band.

### *Pax9* gene cloning, whole-mount in situ hybridization, cryosection, and imaging

Total RNAs were extracted from about 100 embryos (1–3 dpf) using TRIzol reagent (Thermo Fisher, 15596026) according to the manufacturer’s instructions. Reverse transcriptions were performed with the SuperScript® III First-Strand Synthesis System (Thermo Fisher, 18080051) following the manual. The Phusion® High-Fidelity DNA Polymerase master mix (Thermo Scientific, #F530L) was utilized for amplifying the full *pax9* open reading frame (ORF) sequences with primers (forward, 5’-GCCCCCTTGCCACCATGGAGCCAGCCTTTGGGGAGG-3’ and reverse, 5’-CGGCGCGCCCACCCTTTAGAGCTGAAGCCACCAGCGAATGG3-’). The expected-sized PCR products were examined on agarose gels, purified using NucleoSpin Gel and the PCR Clean-up Kit (Takara Bio, 740609.250), and cloned into the pENTR-D vector. *Pax9* ORF sequences were confirmed with nanopore sequencing after cloning.

Riboprobe DNA templates were prepared by plasmid linearization with an endonuclease, NotI (NEB, R0189L), at the 5’ end of the protein-coding region in the pENTR-D vector. The linearized DNA templates were purified prior to in vitro transcription using a NucleoSpin® Gel and PCR Clean-Up kit (Takara Bio, 740609.250). Antisense riboprobes were synthesized by *in vitro* transcription using T7 RNA polymerase (Thermo Scientific, EP0111) and DIG RNA Labeling Mix (Millipore-Sigma, 11277073910) according to the manufacturers’ manuals. All synthesized riboprobes were purified using Sigma Spin post-reaction clean-up columns (Sigma, S5059) and stored at -80°C before use.

Whole-mount *in situ* hybridizations were performed according to our established method, with some modifications ^69^. Briefly, chorions were removed using pronase (Sigma, PRON-RO) treatment before fixation for 0.5-3 dpf fish embryos. All fish embryos were staged and then fixed with 4% PFA (paraformaldehyde) for 1-2 days at 4°C. Color development was carried out in the dark with gentle rocking at room temperature. We closely monitored the color density of each reaction. Once the embryos developed suitable color densities, the reaction was stopped with NTMT washing. The finished samples were imaged immediately or stored in 4% PFA for later imaging. For histological analysis, post-hybridization embryos were equilibrated overnight in 15% sucrose, then in 30% sucrose with 20% gelatin, and finally embedded in 20% gelatin for cryosectioning (10–25 μm) on a cryotome. For imaging, whole-mount embryos were mounted in 3% methylcellulose. Images were acquired using an Axiocam 305 color camera on a Zeiss Stereo Discovery V12 (RRID: SCR_027509) microscope or an Axio Imager 2 compound microscope (RRID: SCR_018876) equipped with Zeiss ZEN Lite software (Ver. 2.3) for image acquisition.

### Heat shock experiments

Fish embryos were collected after breeding and raised in fish system water in a 28 °C incubator. GFP-sorted fish embryos were subjected to a heat shock at 42 °C for 1 hour in egg water in a water bath, as reported at either 24 hpf or 48 hpf ^33,35,36^. After heat shock, embryos were returned to 28 °C in a 10cm-diameter dish in the incubator until 48 hpf. Embryos were then checked and imaged using an Axiocam 305 color camera on a Zeiss Stereo Discovery V12 (RRID: SCR_027509) microscope.

### Fluorescence imaging

For epifluorescence imaging, embryos, larvae, and adult zebrafish were anesthetized with tricaine (MS-222) prior to imaging. For epifluorescence, fish embryos and larvae were mounted in fish system water containing tricaine on a concave slide or in a 3cm-diameter petri dish. Positions were adjusted with a probe needle. Images were acquired using an Axiocam 305 color camera mounted on a Zeiss Stereo Discovery V12 microscope (RRID: SCR_027509), equipped with Zeiss ZEN Lite software (Ver. 2.3) for image acquisition. Fluorescence and brightfield images were captured with appropriate filter sets, and multi-channel images were processed and merged in Fiji (2.16.0/1.54p) and Adobe Photoshop (27.8.0).

For confocal imaging, embryos and larvae were anesthetized with 0.05% tricaine (MS-222) and mounted in 0.6% low-melting-point agarose prepared in fish system water containing tricaine in glass-bottom 3cm diameter dishes and oriented in either left lateral or dorsal position, depending on the tissue being imaged. Fine embryo orientation was adjusted using a probe needle prior to agarose solidification. Confocal fluorescence images were acquired using a spinning-disk confocal system (CrestOptics Cicero) equipped with a Nikon Ti2 inverted microscope and a Lumencor CELESTA light engine. The fluorescence of mNeonGreen, tdTomato, and mCherry was acquired using the corresponding excitation and emission channels. Image acquisition was performed using Nikon NIS-Elements AR software (6.20.02). For z-stack acquisition, the upper and lower boundaries of the target tissue were manually defined, and serial optical sections were collected across the full depth of the region of interest, typically with approximately 100 slices per stack. Z-stack images were processed as maximum-intensity projections in the NIS-Elements software.

### Skeletal preparation, morphological measurement, and fin ray quantification

Larval skeletal preparations were performed using an acid-free two-color cartilage and bone staining protocol adapted from Walker and Kimmel ^70^. Zebrafish larvae and juveniles were euthanized with 0.05% tricaine (MS-222) and fixed in 4% paraformaldehyde (PFA) in PBS at room temperature overnight. Fixed specimens were rinsed and gradient-dehydrated in 25%, 50%, and 75% ethanol for 20 minutes each, with gentle rocking. Cartilage and mineralized bone were stained simultaneously using an acid-free staining solution containing alcian blue 8GX for cartilage and alizarin red S for bone. The staining solution was prepared by mixing the alcian blue solution in 70% ethanol containing 50 mM MgCl₂ with 0.5% alizarin red S immediately before use. Specimens were incubated in staining solution for a few hours to overnight at room temperature with gentle rocking, depending on the stage of staining. After staining, larvae were briefly rinsed in water and bleached in a freshly prepared solution of 1.5% H₂O₂ in 1% KOH to reduce pigmentation. Specimens were then cleared through successive glycerol/KOH solutions (20% glycerol/0.25% KOH, 50% glycerol, 75% glycerol, 100% glycerol) until skeletal structures were clearly visible. Stained specimens were stored in 100% glycerol at 4°C before imaging. Cartilage and bone structures were examined and imaged under an Axiocam 305 color camera mounted on a Zeiss Stereo Discovery V12 microscope (RRID: SCR_027509).

Adult fish were anesthetized with tricaine (Millipore-Sigma, A5040) and imaged under an Axiocam 305 color camera mounted on a Zeiss Stereo Discovery V12 microscope (RRID: SCR_027509). Fin rays were counted manually from fluorescence and brightfield images of caudal, anal, and dorsal fins. FP+ fin rays were identified by fluorescence channel and counted independently. Statistical comparisons between selected pairs of genotypic groups were performed using unpaired two-tailed Welch’s t-tests in GraphPad Prism (10.6.1).

## ACKNOWLEDGMENTS

This research was supported by funding from the National Institute of General Medical Sciences of the National Institutes of Health (R35GM124913) awarded to G.Z. The content is solely the responsibility of the authors and does not necessarily represent the official views of the funding agencies. The authors also thank the Hayward Foundation for its generous support of the laboratory.

