## Supplementary materials for "Spatiotemporal expression of the zebrafish *pax9* gene that is essential for median fin patterning"

Dong and Zhang

**Supplementary materials**

**Figure S1. *Pax9* knockin targeted integration and knockout mutation at the *pax9* locus. A.** Illustration of *pax9* knockin and knockout fish lines based on the Nanopore sequences of the flanking region of the two gRNAs and donor plasmid sequences. **B**. Diagram of *pax9* knockout fish lines, *pu116* and *pu122*. Genomic locus and transcripts information are based on the Ensembl zebrafish genome, GRCz11. Vertical bar indicates the locations of gRNAs.

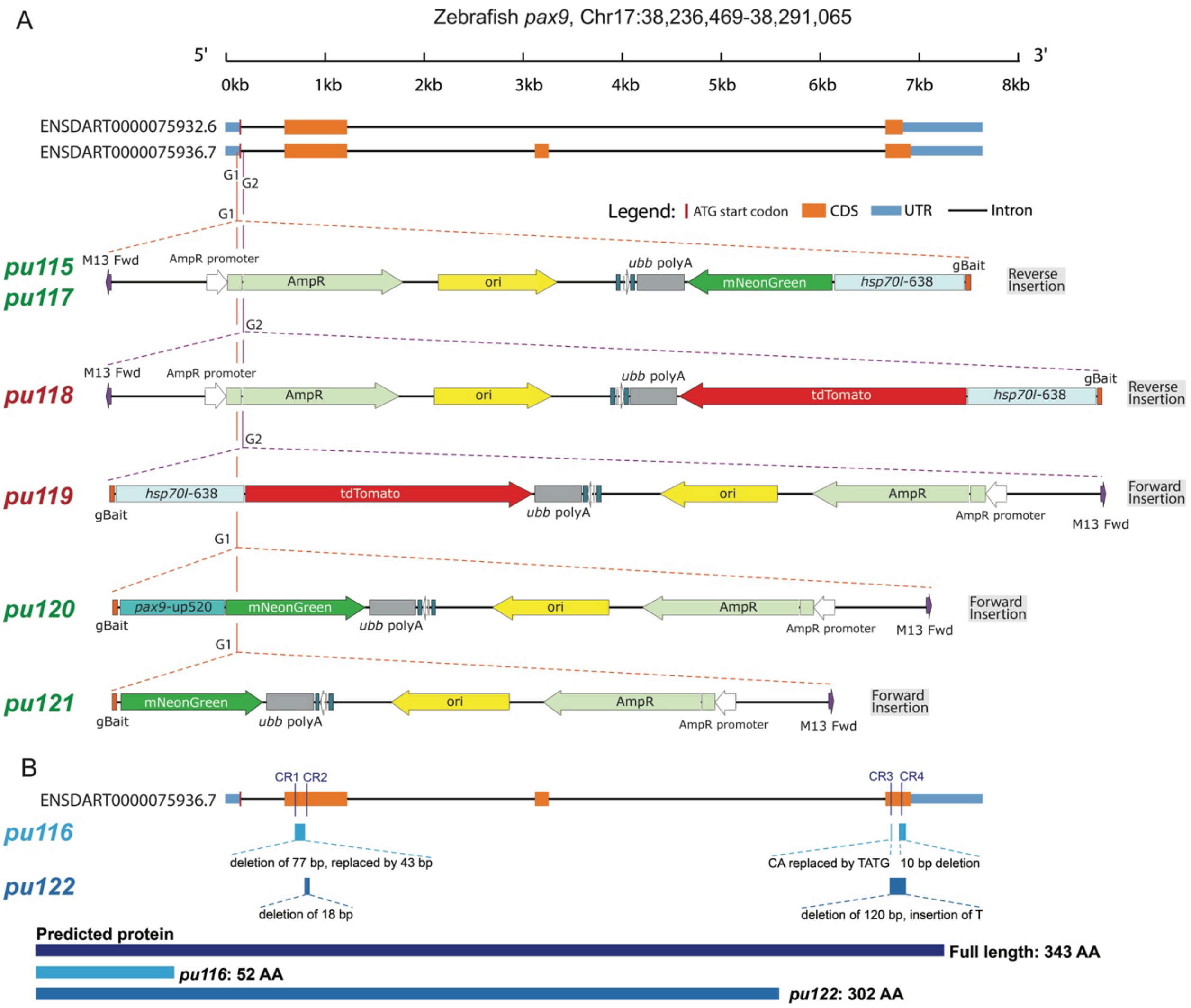


**Figure S2. Increased fluorescent protein-positive fin ray number of transheterozygous knockin median fins. A-E.** Quantification of fluorescent protein-positive (FP+) fin ray number in caudal (total, dorsal lobe, ventral lobe), anal, and dorsal fins of (*pax9*$^{pu119}$, *pax9*$^{pu115}$, and *pax9*$^{pu115/pu119}$). *ns*, not significant. **** indicates significance with p< 0.0001. **F, G.** Two representative anal fins from double knockin fish, *pax9*$^{pu115/pu119}$. FP+ radials and rays are restricted to the anterior region (red arrowheads). **H, I.** Two representative anal fins from knockin fish, *pax9*$^{pu115/pu119}$. Among 18 examined fish, 5 fish showed both anterior (red arrowheads) and posterior-most fluorescence-positive fin rays and radials (yellow arrowheads). **J-K**. Comparison of pectoral and pelvic fin ray numbers of *pax9*$^{pu116/116}$ and non-carrier wildtype fish. 15 fish were counted per genotype for both sides. *ns,* no statistical significance.

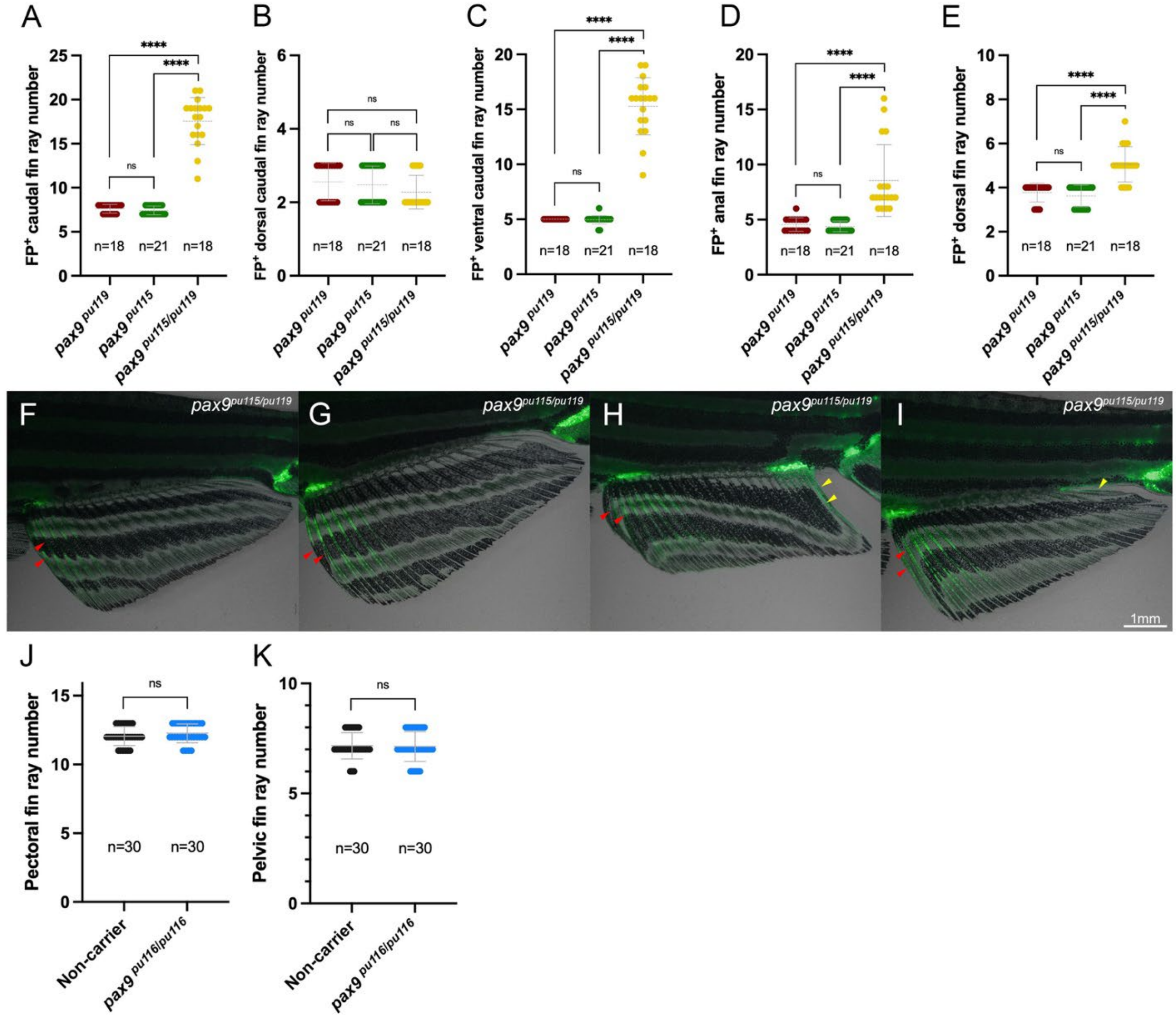


**Table S1. *Pax9* knockin microinjection and screening positive rates.** Four independent microinjections were recorded from $F_0$ to $F_2$ generations.

**Spatiotemporal expression of the zebrafish *pax9* gene, which is essential for median fin patterning**

Dong and Zhang

| F0 injected embryos/fish | | | | | | | F1 fish embryos | | | | | | | F2 fish embryos | | | |
|---|---|---|---|---|---|---|---|---|---|---|---|---|---|---|---|---|---|
| Injected construct | gRNA | Total injected # | Screened # @6dpf | Positive # @6 dpf | Positive screening % | Pos F0% /injection | F0 Founder # | F1 embryos FP | Negative # | Positive # | F1 positive% | Avg pos% | Pos screening % | Negative # | Positive # | Positive% | Avg pos % |
| pUC19-gBait-*hsp70l*-638-mNeonGreen-*ubb* polyA | G1 | 463 | 142 | 21 | 14.79% | 4.53% | #1 Male | negative | 433 | - | - | 24.81% | 40% | | | | 49.06% |
| | | | | | | | #3 Male | negative | 371 | - | - | | | | | | |
| | | | | | | | #4 Male | positive | 58 | 8 | 12.12% | | | | | | |
| | | | | | | | #5 Male | negative | 233 | - | - | | | | | | |
| | | | | | | | #8 Female | positive | 50 | 30 | 37.50% | | | 76 | 81 | 51.59% | |
| pUC19-gBait-*hsp70l*-638-mNeonGreen-*ubb* polyA | G1+G2 | 551 | 167 | 19 | 11.38% | 3.45% | #1 Female | negative | 193 | - | - | 15.14% | 50% | | | | |
| | | | | | | | #2 Female | negative | 286 | - | - | | | | | | |
| | | | | | | | #4 Female | positive | 48 | 16 | 25% | | | | | | |
| | | | | | | | #5 Male | positive | 460 | 6 | 1.29% | | | 165 | 172 | 51.04% | |
| | | | | | | | #6 Female | positive | 55 | 13 | 19.12% | | | 45 | 33 | 42.31% | |
| | | | | | | | #8 Male | incorrect pattern | 78 | - | - | | | | | | |
| pUC19-gBait-*hsp70l*-638-tdTomato-*ubb* polyA | G2 | 569 | 151 | 24 | 15.89% | 4.22% | #1 Female | negative | 268 | - | - | 18.92% | 50% | | | | |
| | | | | | | | #2 Female | positive | 56 | 19 | 25.33% | | | 89 | 83 | 48.25% | |
| | | | | | | | #3 Male | negative | 417 | - | - | | | | | | |
| | | | | | | | #4 Male | positive | 399 | 57 | 12.50% | | | | | | |
| pUC19-gBait-*pax9* up520-mNeonGreen-*ubb* polyA | G1 | 482 | 189 | 29 | 15.34% | 6.02% | #1 Female | negative | 135 | - | - | 7.58% | 37.50% | | | | |
| | | | | | | | #2 Female | negative | 67 | - | - | | | | | | |
| | | | | | | | #3 Female | negative | 13 | - | - | | | | | | |
| | | | | | | | #4 Female | positive | 35 | 2 | 5.41% | | | | | | |
| | | | | | | | #6 Male | positive | 71 | 5 | 6.58% | | | 34 | 37 | 52.11% | |
| | | | | | | | #8 Male | negative | 30 | - | - | | | | | | |
| | | | | | | | #9 Male | positive | 166 | 20 | 10.75% | | | | | | |
| | | | | | | | #10 Male | negative | 39 | - | - | | | | | | |

**Spatiotemporal expression of the zebrafish *pax9* gene, which is essential for median fin patterning**
Dong and Zhang
**Supplementary Protocol**

### Generation of Knockin and Knockout Zebrafish Reporter Lines

This protocol describes the generation of stable zebrafish knockin reporter lines using CRISPR-Cas9-mediated non-homologous end joining (NHEJ) DNA repair mechanism. The overall goal is to integrate a donor plasmid carrying a fluorescent reporter cassette into the 5′ region of an endogenous coding gene of interest, typically near the translation start site. Integration at this position allows endogenous regulatory elements to drive reporter expression while simultaneously disrupting endogenous gene expression, generating a reporter allele that can also function as a loss-of-function allele.

The workflow includes target site selection, guide RNA preparation, donor plasmid construction, injection mix preparation, microinjection into one-cell-stage embryos, $F_0$ fluorescence screening, founder outcrossing, $F_1/F_2$ screening, and molecular validation of targeted integration.

### 1. TARGET SITES SELECTION

#### 1.1 Define the right gene transcript to be labeled.

We generally use the ENSEMBL database to check currently known mRNA transcripts. We will select the common exon with the start codon. If the start-codon-containing exons are different for variable mRNA transcripts, check the literature to determine which one is dominant for your tissue/cell type of interest. Alternatively, these exons can be selected to target them respectively.

For the *pax9* gene, we used the following Ensembl annotation based on the GRCz11 zebrafish genome assembly:

*pax9* Ensembl gene ID: ENSDARG00000053829

- Reference transcript 1: ENSDART00000075932.6
- Reference transcript 2: ENSDART00000075936.7

Both transcripts share the same exon 1 with the start codon (**Fig. S1A**).

#### 1.2 Choose the knockin target region

For reporter knockin at the endogenous locus, select a gRNA target site close to the translation start codon. Ideally, the insertion should occur immediately upstream or downstream of the ATG

- within the 5′ coding region,
- or within the first intron close to the start codon.

For loss-of-function purposes, targeting near the start codon is preferred because donor integration is likely to disrupt the endogenous protein-coding sequence.

It is worth noting that there is likely greater polymorphism and AT-richness in the intron sequences. Moreover, the variable zebrafish strains used across different labs could be diverse. So the standard sequences from ENSEMBL may differ from those of your zebrafish strain. Ideally, it is better to sequence the region that will be used for gRNA design.

#### 1.3 Design gRNAs and PCR primers

Candidate gRNAs and flanking PCR primers can be designed using CHOPCHOP or other CRISPR design tools.

Recommended criteria:

- SpCas9-compatible NGG PAM
- high on-target score
- low predicted off-target score
- located close to the ATG or intended insertion site
- avoids common SNPs if strain-specific genome information is available
- preferably produces a cut site compatible with donor insertion and downstream genotyping

For *pax9*, the following two gRNAs were designed:
G1: 5'-GACTCGGAACAGGTCAGAAT-3', targeting *pax9* exon 1 immediately upstream of the ATG.
G2: 5'-ATGCATGCCCGCTCCCATGT-3', targeting *pax9* intron 1 immediately downstream of the ATG.

### 2. GRNA PREPARATION

#### 2.1 Design gRNA template oligonucleotides

We use the Gagnon strategy for gRNA preparation [1]. For each target DNA template, there are 3 elements:

1. an *in vitro* transcription promoter sequence T7 or Sp6;
2. a target-specific spacer sequence;
3. a constant overlap sequence for sgRNA scaffold template generation.

Because of the cost of longer oligonucleotide synthesis, we synthesize two oligonucleotides and then anneal and ligate them into a single gRNA template. We generally choose T7 promoter sequence due to its higher efficiency. As the G is critical for T7 promoter efficiency, the first nucleotide may be changed to G if the target spacer does not begin with G.
For *pax9*, we selected G1 and G2 gRNA sequences:
G1 GACTCGGAACAGGTCAGAAT—AGG (PAM)
G2 ATGCATGCCCGCTCCCATGT—GGG (PAM)

Corresponding 60-mer oligonucleotides
T7pax9-G1 5'-
TAATACGACTCACTATAGACTCGGAACAGGTCAGAATGTTTTAGAGCTAGAAATAGCAAG-3'
T7pax9-G2 5'-
TAATACGACTCACTATAGTGCATGCCCGCTCCCATGTGTTTTAGAGCTAGAAATAGCAAG-3'

Shared constant oligonucleotides:
5'-
AAAAGCACCGACTCGGTGCCACTTTTTCAAGTTGATAACGGACTAGCCTTATTTTAACTTGCT ATTTCTAGCTCTAAAAC-3'

Blue = T7 RNA polymerase binding and recognition sequence
Red = gRNA sequence
Purple = constant oligo overlapping sequence

#### 2.2 Anneal gene-specific and constant oligonucleotides

Prepare the annealing reaction in a 200ul PCR tube.

| Component | Volume |
|---|---|
| **Gene-specific oligo, 100 µM** | 2 µL |
| **Constant oligo, 100 µM** | 2 µL |
| **Nuclease-free water** | 16 µL |
| **Total** | 20 µL |

Then incubate the samples in a Thermocycler and run the following program.

1. 95°C for 10 min
2. cool from 95°C to 85°C at 0.3 °C/s
3. cool from 85°C to 25°C at 0.1 °C/s
4. hold at 12°C

#### 2.3 Fill-in and repair reaction

Prepare the fill-in reaction to generate a double-stranded DNA template for *in vitro* transcription.

| Component | Volume |
|---|---|
| **Annealed oligo reaction** | 20 µL |
| **dNTPs (10 mM)** | 5 µL |
| **10× NEB Buffer 2** | 4 µL |
| **100x NEB BSA** | 0.4 µL |
| **T4 DNA polymerase (NEB, M0203S )** | 1 µL |
| **Nuclease-free water** | 9.6 µL |
| **Total** | 40 µL |

Incubate at 12°C for 20 minutes.

Optional: Check template quality by running 1 µL of the fill-in reaction on a 1-2% agarose gel. One ~120 bp band should be visible. The fill-in product was purified before being used as the DNA template for in vitro transcription using the NucleoSpin™ Gel and PCR Cleanup kit (Takara Bio, 740880.250).

#### 2.4 *In vitro* transcription of gRNA

Set up the transcription reaction using a T7 (or SP6) in vitro transcription kit (HiScribe® T7 High Yield RNA Synthesis Kit, E2040S).

| Component | Volume |
|---|---|
| **ATP** | 1.5 µL |
| **GTP** | 1.5 µL |

| | |
|---|---|
| **CTP** | 1.5 µL |
| **UTP** | 1.5 µL |
| **10× transcription buffer** | 1.5 µL |
| **T7/SP6 enzyme mix** | 2 µL |
| **DNA template (1000 ng DNA)** | X µL |
| **Nuclease-free water** | (fill up to 20) µL |
| **Total** | 20 µL |

Incubate at 37°C overnight.
After transcription, add DNase to remove template DNA:

| **Component** | **Volume** |
|---|---|
| **Transcription reaction** | 20 µL |
| **DNase** | 1 µL |

Incubate at 37°C for 30-60 min.

#### 2.5 gRNA purification

Purify gRNA using column-based RNA cleanup (alternative: ethanol/ammonium acetate precipitation) gRNA column purification: Monarch® Spin RNA Cleanup Kit (NEB, T2050L).

Example column purification workflow:

1. Adjust the transcription reaction volume to 50 µL by adding 30 µL RNase-free water.
2. Add 100 µL RNA Binding Buffer. Mix by vortexing and briefly centrifuge.
3. Add 150 µL ethanol. Mix by vortexing and briefly centrifuge.
4. Load the mixture onto the column. If precipitate remains on the tube wall after brief centrifugation, resuspend it by pipetting and transfer the entire mixture to the column.
5. Centrifuge at 14,800 rpm (maximum speed) for 1 min.
6. Wash the column twice with 500 µL RNA Wash Buffer. First wash at 3,000 rpm, and second wash at 14,800 rpm.
7. After the final wash, centrifuge the empty column for 3 min at full speed to remove residual ethanol.
8. Elute RNA in 50 µL nuclease-free water or elution buffer. For a higher yield, the eluate can be reapplied to the column for a second elution (50 µL + 50 µL).

#### 2.6 gRNA quality control and storage

Measure RNA concentration using Nanodrop or an equivalent method.
Recommended QC:

- concentration: >500 ng/µL
- A260/280: ~2.0
- A260/230: ~2.0-2.2
- optional gel electrophoresis check for RNA integrity

Prepare aliquots or a 500 ng/µL working solution to avoid repeated freeze-thaw cycles.
Store gRNA aliquots at −80°C.

### 3. *IN VIVO* GRNA EFFICIENCY TEST

The cutting efficiency of the target-gene gRNA is a major factor affecting knockin efficiency. Based on our experience, *in vitro* and *in vivo* gRNA-cutting efficiencies may differ markedly. Therefore, before performing the knockin experiment, we generally tested each candidate gRNA *in vivo* by microinjection followed by a T7E1 assay. This step helps identify gRNAs with strong activity in zebrafish embryos and increases the likelihood of recovering fluorescently positive knockin founders during subsequent screening.

#### 3.1 Prepare gRNA efficiency test injection solution

Prepare the pre-Cas9 injection mix fresh on the day of injection when possible. Alternatively, aliquots can be prepared in advance, stored at -80°C, and thawed on ice before use.

Example preparation of pre-Cas9 injection mix for three injections:

| Component | Stock concentration | Concentration in pre-Cas9 mix | Volume |
|---|---|---|---|
| **Target gene gRNA** | 500 ng/µL | 50 ng/µL | 0.9 µL |
| **Phenol red** | - | - | 3 µL |
| **Nuclease-free water** | - | - | 5.1 µL |
| **Total** | - | - | 9 µL |

Mix thoroughly and briefly centrifuge. Aliquot into 3 µL per tube and keep on ice if used immediately, or store at -80°C for later use.

#### 3.2 Microinjection

The injection process was described in our published protocol [2]. For detailed injection setup and procedures, see Sections 5 and 6.

Immediately before injection, thaw one 3 µL aliquot of pre-Cas9 injection mix on ice and add 1 µL Cas9 protein (1 µg/µL, 6 µM). Mix gently by pipetting, briefly centrifuge, then incubate at room temperature for 5 min to allow formation of the Cas9-gRNA ribonucleoprotein complex. Keep the injection mix on ice until use.

Back-load the injection needle with approximately 1 µL of injection mix, then break the needle tip with fine forceps under a stereomicroscope. Collect zebrafish embryos immediately after spawning, sort healthy one-cell-stage embryos, and place embryos on agarose injection molds in fish water or E3 medium. Inject approximately 2 nL of injection solution into the cell of one-cell-stage embryos. For a gRNA efficiency test, 50-100 injected embryos are usually sufficient.
After injection, transfer embryos to clean fish water or E3 medium and incubate at 28.5°C.

#### 3.3 Genomic DNA collection

Embryo genomic DNA can be collected at 1-2 dpf. Remove chorions either manually using forceps or by pronase digestion. For pronase digestion, add 20 µL of pronase to the petri dish and incubate the embryos at room temperature until the chorions are removed; then rinse the embryos thoroughly with fish water.

Remove deformed fish embryos and pool 15-20 healthy embryos in a 1.5 mL microcentrifuge tube or 200 μL PCR tube. Remove excess liquid and add 100 μL of 50 mM NaOH. Incubate at 96°C for 1 h to lyse embryos. Then neutralize the lysate by adding 10 μL of 1 M Tris-HCl (pH 7.9). Vortex briefly and centrifuge to pellet debris. Use the supernatant as a genomic DNA template for PCR.

#### 3.4 T7E1 assay and quantification

Design PCR primers flanking the gRNA target site. An expected PCR product size of approximately 200-400 bp is recommended for clear visualization of T7E1 digestion products.

1) PCR amplification of the regions around the gRNA targets.

Set up the following PCR reaction:

| Component | Volume |
|---|---|
| **10x Taq buffer** | 5 μL |
| **50 mM MgCl2** | 3.6 μL |
| **10 mM dNTPs** | 1 μL |
| **Forward primer** | 1 μL |
| **Reverse primer** | 1 μL |
| **DNA template** | 2 μL |
| **Taq Polymerase** | 1 μL |
| **Water** | 35.5 μL |
| **Total** | 50 μL |

Incubate PCR samples in a thermocycler with the following program:

| Step | Temperature | Time | Cycles |
|---|---|---|---|
| **Initial denaturation** | 95°C | 1min | 1 |
| **Denaturation** | 95°C | 30s | 35x |
| **Annealing** | 60°C | 30s | 35x |
| **Extension** | 72°C | 1min | 35x |
| **Final extension** | 72°C | 5min | 1 |

2) PCR product DNA preparation by denaturing and renaturing

This denaturation-renaturation step allows heteroduplex formation between wild-type and mutant PCR products.

Set up the following reaction:

| Component | Volume |
|---|---|
| **PCR product** | 18 μL |
| **NEB Buffer 2** | 2 μL |
| **Total** | 20 μL |

Incubate PCR samples in a thermocycler with the following program:

1. 95°C for 10 min
2. cool from 95°C to 85°C at 0.3 °C/s
3. cool from 85°C to 25°C at 0.1 °C/s
4. hold at 12°C

3) T7 Endonuclease I digestion:

The T7 Endonuclease I (T7E1) recognizes and cleaves mismatched heteroduplex DNA. It can therefore be used to estimate CRISPR-Cas9-induced mutation frequency at the target locus.

Set up the following reaction:

| **Component** | **Volume** |
|---|---|
| **Denatured/renatured PCR product** | 20 μL |
| **NEB Buffer 2** | 0.2 μL |
| **T7E1 enzyme (NEB, M0302S)** | 0.5 μL |
| **H2O** | 1.3 μL |
| **Total** | 22 μL |

Incubate at 37°C for 1h in a thermocycler. Then immediately check the results on an agarose gel or keep the samples frozen to prevent further digestion.

4) Agarose gel electrophoresis and quantification

Run both the undigested PCR product and the T7E1-digested product on a 1.5-2% agarose gel. Image the gel and quantify band intensity using ImageJ greyscale analysis.

Cutting efficiency can be estimated using the following formula:

$T7E1\ efficiency = \frac{WT\ band - T7E1\ remaining\ band}{WT\ band}$.

For the two *pax9* gRNAs tested in this protocol, G1 showed an estimated cutting efficiency of 82.92%, and G2 showed an estimated cutting efficiency of 92.42%.

Based on our experience, we recommend prioritizing gRNAs with estimated *in vivo* cutting efficiency above 80% for knockin injections.

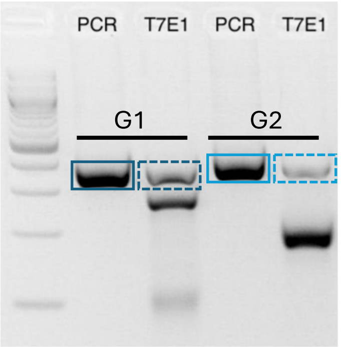


**Example of *pax9* gRNA T7E1 efficiency test.** Solid line boxes indicate PCR band; dotted line boxes indicate T7E1 cleavage remaining band.

### 4. DONOR PLASMID DESIGN AND PREPARATION

#### 4.1 Donor plasmid design

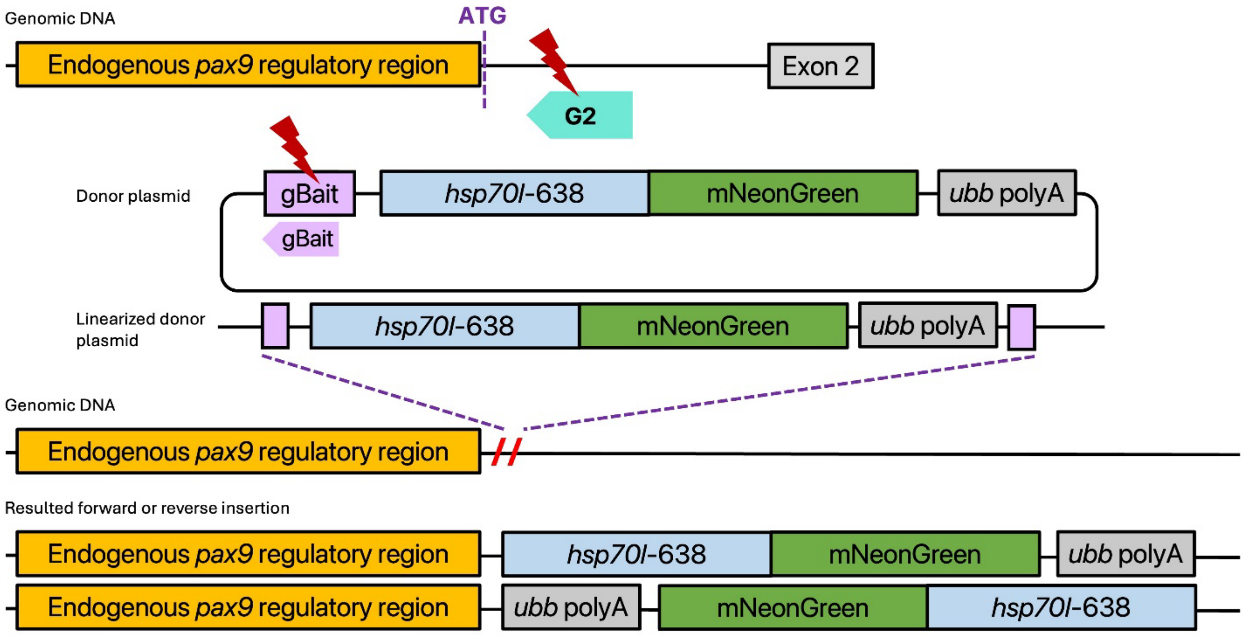


**Illustration of *the pax9 knock-in* strategy.** *Pax9* gRNA and gBait gRNA allow cleavage of the endogenous *pax9* locus and donor plasmid. So the linearized donor cassette can be integrated in either forward or reverse orientation.

The donor plasmid contains a reporter or driver cassette and a gBait sequence that can be cleaved by the gBait gRNA during injection. After Cas9/gBait-mediated cleavage, the linearized donor can integrate into the target genomic locus through non-homologous end joining (NHEJ). Because NHEJ-mediated integration can occur in either orientation, downstream junction PCR and sequencing are required to determine the insertion orientation and validate the allele.

General donor architecture:
Bait sequence -- promoter/enhancer element -- reporter/driver cassette -- *ubb* polyA -- plasmid backbone
For the *pax9* knockin fish lines described here, example donor plasmids include:

pUC19-gBait-*hsp70l*-638-mNeonGreen-*ubb* polyA
pUC19-gBait-*hsp70l*-638-tdTomato-*ubb* polyA

#### 4.2 Preparation of high-quality donor plasmid DNA for injection

High-quality plasmid DNA is important for embryo survival and knockin efficiency. We choose TakaraBio NucleoBond® Xtra Midi for this purpose. DNA quality should meet the following standards.

| **Parameter** | **Recommended range** |
|---|---|
| **Concentration** | >500 ng/μL |
| **A260/280** | ~1.8 |
| **A260/230** | >1.8 if possible |

| | |
|---|---|
| **RNA contamination** | minimal |
| **Genomic DNA contamination** | minimal |

#### 4.3 Diagnostic digestion

After midi-prep high-quality plasmid preparation, perform a diagnostic restricted endonuclease digestion and run the product on a 0.8% agarose gel to confirm plasmid identity. The correct plasmid can be stored at -20°C for a long time.

### 5. INJECTION SOLUTION PREPARATION

#### 5.1 Injection mix components

Prepare the injection mix fresh on the day of injection when possible. Alternatively, an aliquot of the injection mix without Cas9 protein (PNA BIO CP01-50) can be prepared in advance, stored at −80°C, and thawed on ice immediately before use. All tubes and tips should be RNase-free.

Example of injection solution preparation (3 injections):

| Component | Stock concentration | Final cc (pre-Cas9 mix) | Volume |
|---|---|---|---|
| **Target gene gRNA** | 500 ng/μL | 50 ng/μL | 0.9 μL |
| **gBait gRNA** | 500 ng/μL | 50 ng/μL | 0.9 μL |
| **Donor plasmid*** | 300 ng/μL | 30 ng/μL | 0.9 μL |
| **Phenol red** | - | - | 3 μL |
| **Nuclease-free water** | - | - | 3.3 μL |
| **Total** | - | - | 9 μL |

*Volume of donor plasmid is determined by plasmid concentration.
Mix thoroughly and briefly centrifuge. Aliquot into 3 μL per tube and keep on ice if used immediately, or store at -80°C for later use.

#### 5.2 Complex formation

Immediately before injection:

1. Thaw one 3 μL aliquot of injection mix on ice and add 1 μL Cas9 protein (1μg/μL, 6μM, PNA BIO CP01-50) into 3 μL injection mix. Mix gently by pipetting and briefly centrifuge. (After addition of Cas9 protein at a 3:1 mix-to-Cas9 ratio, all non-Cas9 components are diluted by 25%)
2. Incubate at room temperature for 5 minutes to allow ribonucleoprotein complex formation.
3. Keep the injection mix on ice until use.

#### 5.3 Needle loading

Injection needles were prepared by using the Sutter Instrument Model P-1000 (Flaming/Brown micropipette puller) to pull injection needles (World Precision Instruments, Thin wall glass capillaries, 4 Inch 1.0 OD, TW100F-4). The following parameters were used in the Sutter Instrument Model P-1000 machine:

| Parameter | Setting |
|---|---|
| **Heat** | 561 |
| **Pull** | 90 |
| **Velocity** | 100 |
| **Time** | 250 |
| **Pressure** | 450 |

Back-load the injection needle with 1-1.5 µL injection mix using a 20 µL microloader tip (Calibre Scientific Microloader, Catalog # 930001007). Break the needle tip using fine forceps under a stereomicroscope. Before injection, calibrate the injection volume by ejecting droplets into mineral oil and adjusting injection pressure/time until the desired volume is reached. For this protocol, approximately 2 nL of injection solution was injected per embryo.

#### 6. Zebrafish embryo collection and microinjection

The injection process was described in our published protocol [2].

#### 6.1 Breeding setup

Set up adult zebrafish breeders the evening before injection.
Example setup:

- Wild type strain: TAB
- Fish number: 3 females + 3 males
- number of tanks: 4
- Put dividers to separate males and females in breeding tanks.

On the morning of injection day, remove dividers from one or two tanks at a time. Collect embryos immediately after spawning, ideally within 5–10 min. If additional embryos are needed, replace the dividers and repeat embryo collection from additional tanks.

#### 6.2 Embryo preparation

Sort healthy one-cell-stage embryos and remove unfertilized, abnormal, opaque, or white-appearing embryos. Embryos are injected through the chorion; therefore, dechorionation is not required before injection. Place embryos on agarose injection molds/injection plates in fish water or E3 medium. Align fish embryos along the lanes on the mold.

#### 6.3 Microinjection

Inject ~2 nL injection solution into the cell of one-cell-stage embryos.
Important points:

- For efficient knockin, injections should be performed at the early one-cell stage.
- To maximize the number of embryos injected at the early one-cell stage, collect and inject embryos in smaller batches. For example, collect approximately 150 embryos, complete

injection within 30 min, and repeat this process 2 or 3 times, rather than collecting 500 embryos and injecting for 2 h.
- Inject into the cell rather than the yolk whenever possible. At the early one-cell stage, the cell appears flatter on top of the yolk.
- Avoid damaging the cell or yolk by injecting excessive volume.
- After injection, transfer embryos to clean fish water and incubate at 28.5°C.
- Record the number of injected embryos, injection date, gRNA, donor plasmid, Cas9 batch, and injection mix details.

#### 6.4 Post-injection care

Incubate injected embryos at 28.5°C. At the end of the injection day, remove unfertilized or damaged embryos to maintain water quality. In the following days, check embryo survival and gross morphology frequently.
If the injection is highly toxic, optimize the injection process by adjusting:
- Cas9 concentration
- target gRNA and gBait gRNA concentration
- donor plasmid concentration
- injection volume
- needle size
- whether the injection process causes excessive damage to the cell or yolk

### 7. $F_0$ EMBRYO AND LARVAL SCREENING

#### 7.1 Screening stage

Screen the injected $F_0$ embryos or larvae at stages when endogenous target gene expression is expected. This stage varies by gene. Whole-mount *in situ* hybridization or other published expression data can be used as a reference for the target gene's spatiotemporal expression pattern.
For $F_0$ injected embryos, fluorescent protein signals are often weak, mosaic, and sporadic. Candidate $F_0$ larvae should therefore be selected based on fluorescence in expected endogenous expression domains rather than overall brightness alone. For our *pax9* reporter screening, potential founder larvae were selected by identifying pharyngeal arch-specific fluorescence at 3-4 dpf.

#### 7.2 Fluorescence screening

Screen $F_0$ embryos/larvae using a fluorescence stereomicroscope.
Screening categories:

| Category | Description | Raise? |
|---|---|---|
| No fluorescence | no detectable reporter signal | no |
| Weak nonspecific fluorescence | scattered or ectopic signal | no |
| Strong broad ectopic fluorescence | widespread fluorescence inconsistent with expected expression | no |

| | | |
|---|---|---|
| Mosaic expected-domain fluorescence | signal in the expected tissue domain | yes |
| Strong expected-domain fluorescence | clear signal in the endogenous domain | yes |
| Severe abnormal morphology | developmental defects | no |

For knockin injection optimization, record the number of embryos in each category. Knockin efficiency may vary across injection batches and be affected by injection performance, gRNA cleavage efficiency, donor plasmid quality, Cas9 activity, and embryo quality. In some cases, the target gene may have low or restricted expression. Confocal microscopy may be used when fluorescence signals are too weak to be reliably identified with a stereomicroscope.

#### 7.3 Candidate $F_0$ selection

Select $F_0$ larvae showing reporter expression consistent with the expected endogenous pattern. Avoid selecting larvae with only broad nonspecific fluorescence. Raise selected $F_0$ larvae to adulthood under standard zebrafish husbandry conditions.

### 8. GERMLINE TRANSMISSION SCREENING

#### 8.1 $F_0$ outcrossing

When $F_0$ fish reach sexual maturity, outcross each candidate $F_0$ adult to wild-type fish. Collect $F_1$ embryos and screen for fluorescence at the expected developmental stage.

Example recording table:

| $F_0$ ID | Sex | Cross date | # of $F_1$ embryos | Number FP+ | Transmission rate | Pattern |
|---|---|---|---|---|---|---|
| ***pu115* #1** | Female | 2026.4.15 | 200 | 10 | 5% | sclerotome@20hpf, pharyngeal arch@3 dpf |

Transmission rate = fluorescent protein positive embryos / total embryos screened × 100%

#### 8.2 Criteria for positive founders

A founder is considered germline positive if its $F_1$ embryos show:

- fluorescent reporter expression in expected endogenous domains
- reproducible expression among multiple embryos
- survival compatible with raising
- preferably a Mendelian transmission pattern in later generations

Raise positive $F_1$ embryos to adulthood. The $F_1$-positive rate from $F_0$ knockin founders may vary substantially among individuals because $F_0$ animals are mosaic. In practice, if no fluorescent-positive embryos are observed after screening approximately 200 $F_1$ embryos from an outcross, the $F_0$ adult is considered unlikely to be a germline transmitter and is typically classified as negative.

### 9. ESTABLISHMENT OF $F_2$ STABLE LINES

#### 9.1 $F_2$ fish generation and screening

Outcross $F_1$ adults to wild-type fish. Screen $F_2$ embryos for fluorescence at the expected developmental stage. If the line carries a single-locus heterozygous insertion, the expected fluorescent-positive rate from an outcross should approach approximately 50%, although the observed rate may vary depending on viability, scoring stage, fluorescence strength, and insertion complexity. Stable lines should be maintained by outcrossing fluorescent-positive carriers to wild-type fish. Try to establish more than 2 founders to compare the consistency of knockin.

#### 9.2 Line naming

Assign stable line names after confirmation of germline transmission and molecular validation. Follow the ZFIN Zebrafish Nomenclature Conventions.

Recommended line record:

| Stable line | Founder ID | Donor construct | Reporter | Target gRNA | Insertion orientation | F2 FP+ rate | Molecular validation |
|---|---|---|---|---|---|---|---|
| ***pu115*** | #1 female | pUC19-gBait-*hsp70l*-638-mNeonGreen-*ubb* polyA | mNeonGreen | G1 | reverse | 50% | PCR and sequencing done |

For each stable line, record the line name, donor construct, reporter, target gRNA, insertion orientation, fluorescence pattern, and molecular validation status.

### 10. MOLECULAR VALIDATION OF KNOCKIN ALLELES

#### 10.1 Genomic DNA preparation

Genomic DNA was prepared from caudal fin clips of adult fluorescent-positive fish from stable knockin lines.

For adult fin clips:

1. Anesthetize fish with 0.05% tricaine (MS-222).
2. Clip a small piece of a caudal fin using a sterile blade. Fish were returned to the fish system water for recovery after fin clipping.
3. Fin tissue was transferred directly into lysis buffer, and genomic DNA was extracted using the NucleoSpin® DNA RapidLyse kit according to the manufacturer's instructions.

#### 10.2 Junction PCR

Junction PCR was performed to confirm targeted integration at the endogenous locus and to determine donor cassette orientation. Primers were designed so that one primer annealed to the genomic sequence outside the CRISPR/Cas9 cut site and the paired primer annealed within the donor cassette.

Design PCR primers to test integration at the endogenous locus:

1. 5′ junction PCR: one primer outside the genomic cut site + one primer inside donor cassette
2. 3′ junction PCR: one primer inside donor cassette + one primer outside the genomic cut site
3. Orientation-specific primer combinations were used to distinguish forward and reverse donor cassette insertion, as illustrated in the diagram.

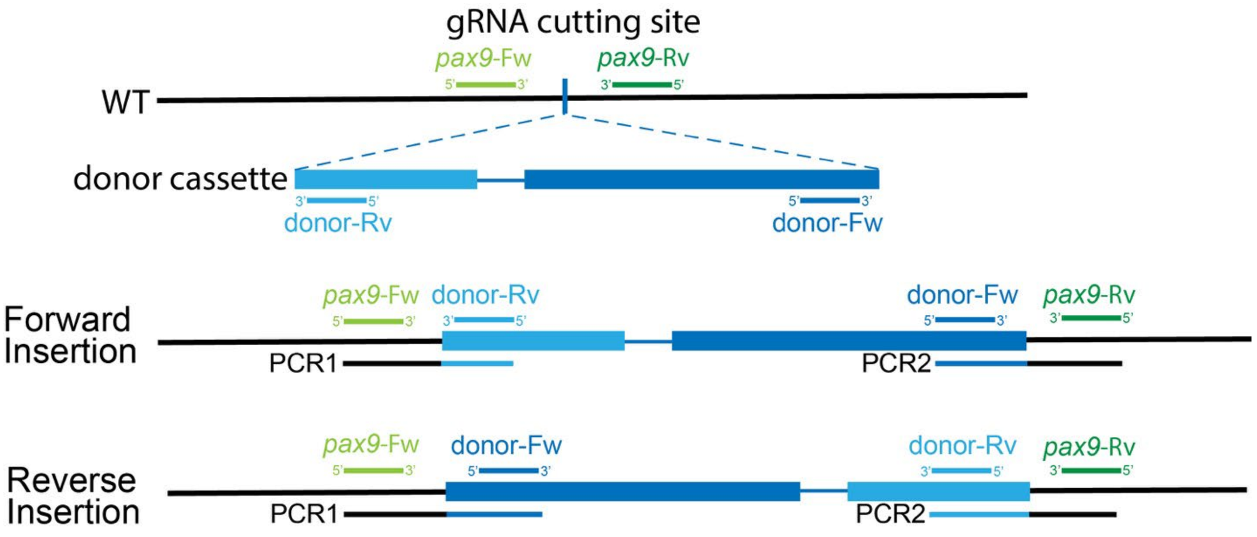


PCR reactions were performed using the HyperFusion (APExBIO, K1032 ) or Phusion (NEB, M0530S) high-fidelity DNA polymerase kit.

- Test insertion orientation

Prepare the PCR reaction as follows.

| Component | Volume |
|---|---|
| **5× HyperFusion buffer** | 4 μL |
| **2.5 mM dNTPs** | 1.6 μL |
| **Forward primer** | 0.5 μL |
| **Reverse primer** | 0.5 μL |
| **DNA template** | 2 μL |
| **Polymerase** | 0.4 μL |
| **Water** | 11 μL |
| **Total** | 20 μL |

Run the samples on a thermocycler with the following PCR program.

| Step | Temperature | Time | Cycles |
|---|---|---|---|
| **Initial denaturation** | 98°C | 1min | 1 |
| **Denaturation** | 98°C | 15s | 35x |
| **Annealing** | 58°C | 15s | 35x |
| **Extension** | 72°C | 30s/kb | 35x |

| | | | |
|---|---|---|---|
| **Final extension** | 72°C | 5min | 1 |

PCR products were analyzed on a 1% agarose gel to determine whether the expected orientation-specific bands were present.

#### 10.3 Cloning and sequencing of insertion junctions

Once insertions and their directions were verified by PCR, proceed with gene cloning and sequencing. For gel purification and downstream cloning, larger 50 µL PCR reactions were prepared using the same primer combinations and cycling conditions:

| Component | Volume |
|---|---|
| **5× HyperFusion buffer** | 10 µL |
| **2.5 mM dNTPs** | 4 µL |
| **Forward primer** | 1 µL |
| **Reverse primer** | 1 µL |
| **DNA template** | 2 µL |
| **Polymerase** | 1 µL |
| **Water** | 31 µL |
| **Total** | 50 µL |

PCR products were separated on a 0.8% agarose gel, and target bands of the expected size were excised and purified using the NucleoSpin™ Gel and PCR Cleanup kit (Takara Bio, 740880.250 ) according to the manufacturer's instructions.

Purified junction PCR products were then cloned into the pJET1.2 vector using CloneJET PCR Cloning Kit (K1232). After transformation and colony selection, plasmid DNA was prepared by miniprep. Candidate clones were identified by BglII restriction endonuclease digestion to verify the presence of an insert of the expected size. If needed, additional restriction enzymes were selected based on the pJET plasmid map and the predicted PCR product sequence to determine insert orientation within the vector.

Two plasmids with the expected insert size and sufficient DNA concentration were submitted for Nanopore or Sanger sequencing. Ideally, the plasmid should have a concentration> 100 ng/µL to ensure sequencing quality and accuracy. Sequence analysis was used to determine the precise genomic insertion site, donor cassette orientation, and local indels at the insertion junction. Knockin alleles were considered validated when junction PCR and sequencing confirmed that the donor cassette was inserted at the intended genomic locus and identified the exact sequences where the donor cassette joined the genomic DNA.
